# Biogeography shapes the TE landscape of *Drosophila melanogaster*

**DOI:** 10.1101/2025.05.22.655554

**Authors:** Riccardo Pianezza, Robert Kofler

## Abstract

The abundance and composition of transposable elements (TEs) varies widely across species, yet the evolutionary forces shaping this diversity remain poorly understood. Using 285 recently published genomes from drosophilid species, we investigated the evolutionary origins of the ≈130 TE families present in *D. melanogaster* and found that 79 were exchanged via horizontal transposon transfer (HTT) with other drosophilids.

Most HTT events involved closely related species such as *D. simulans, D. mauritiana*, and *D. teissieri*, although transfers from more distantly related taxa were also observed. Notably, *D. melanogaster* appears to be a net recipient of HTTs, acquiring about three times as many TEs as it donated.

Geographic patterns reveal that most HTTs involved Afrotropical species, reflecting *D. melanogaster* ‘s ancestral range, with fewer involving species from the Neotropics, a region which *D. melanogaster* invaded only ≈200 years ago. Despite colonizing the Nearctic, Australasian, and Palearctic regions between 200–2000 years ago, we found no evidence of HTT with species from those areas.

Nonetheless, an analysis of drosophilids from each biogeographic realm shows that HTT is widespread in each realm, with 3–55% of the genome in each species derived from HTT. Strikingly, a considerable portion of the genome is shared among all species inhabiting the same realm —regardless of phylogenetic distance—indicating that geographic overlap, rather than shared ancestry, is a primary driver of TE composition. These findings highlight biogeography as a major force shaping the TE landscape and underscore the importance of ecological interactions in genome evolution.

## 1 Introduction

Transposable elements (TEs) are selfish genetic elements that can replicate within a host genome. They are present in all sequenced eukaryotic genomes investigated so far [Wicker et al., 2007]. Two main classes of TEs can be distinguished: DNA transposons, which replicate with a ‘cut and paste’ mechanism, and retrotransposons, which utilize a ‘copy and paste’ mechanism [Finnegan, 1989, Wicker et al., 2007]. For retrotransposons, we can further distinguish between long terminal repeat (LTR) and non-LTR transposons. The genomic proportions of TEs ranges from 2% in the Japanese pufferfish [Aparicio et al., 2002] to 85% in maize [Stitzer et al., 2021]. In humans, TEs account for 55% of the genome [Nurk et al., 2022]. To prevent the spread of TEs, host organisms have evolved sophisticated defense mechanisms, which frequently involve small RNAs [Sarkies et al., 2015]. In *Drosophila*, the host defense is based on piRNAs: small RNAs ranging in size from 23-29 nt that mediate the silencing of TEs at transcriptional and post-transcriptional levels [Brennecke et al., 2007, Czech and Hannon, 2016]. Over time, inactive TEs gradually accumulate mutations, reducing their ability to mobilize. Eventually, due to this gradual erosion, a population may have no functional copies of a TE remaining, resulting in the death of the TE family [Blumenstiel, 2019, Schaack et al., 2010]. TEs can avoid this evolutionary demise through two strategies. First, TEs that can maintain at least some functional copies over evolutionary timescales may be vertically transmitted. A weak residual activity, despite an active host defense, could help maintain functional copies of a TE family. In particular, non-LTRs transposons may be frequently vertically transmitted [Zhang et al., 2020, Schaack et al., 2010, Loreto et al., 2008]. Another strategy that may facilitate the long-term persistence of a TE is horizontal transfer (HT) to a species that lacks the TE [Blumenstiel, 2019, Schaack et al., 2010]. This HT may trigger a burst of TE activity in the novel host, that stops when the TE is again silenced by the host defense. The horizontal transfer of TEs (HTT) is abundant in bacteria, but it may also be common in eukaryotes [Keeling and Palmer, 2008]; An investigation of 195 insect genomes revealed 2248 HTT in insects [Peccoud et al., 2017]. Due to methodological limitations, the authors believe that the true extent of HTT has been underestimated by several orders of magnitude.. This was strikingly confirmed by recent studies that identified 11 TE invasions, likely triggered by HTT, in *D. melanogaster* over the past 200 years [Scarpa et al., 2023, Schwarz et al., 2021, Bucheton et al., 1992, Periquet et al., 1989, Bonnivard et al., 2000, Kidwell, 1983, Anxolabéhère et al., 1988, Daniels et al., 1990, Pianezza et al., 2023, 2024]. These invasions increased the *D. melanogaster* genome by 1%, representing a massive genome change in a short period of time. For 8 of these 11 TEs, the donor species is likely *D. simulans*, a cosmopolitan species closely related to *D. melanogaster* [Capy and Gibert, 2004]. *The remaining 3 TEs were likely acquired from a Neotropical Drosophila* species [Daniels et al., 1990, Pianezza et al., 2023, 2024]. Importantly, *D. melanogaster* extended its range into the Neotropics during the last 200 years [Chen et al., 2024]. Therefore, the range expansion into the Neotropics is indirectly responsible for the HTT of these three TEs. In general, the biogeographic history of a species could be a key factor in shaping the horizontal exchange of TEs between species [Peccoud et al., 2017].

**Figure 1:**
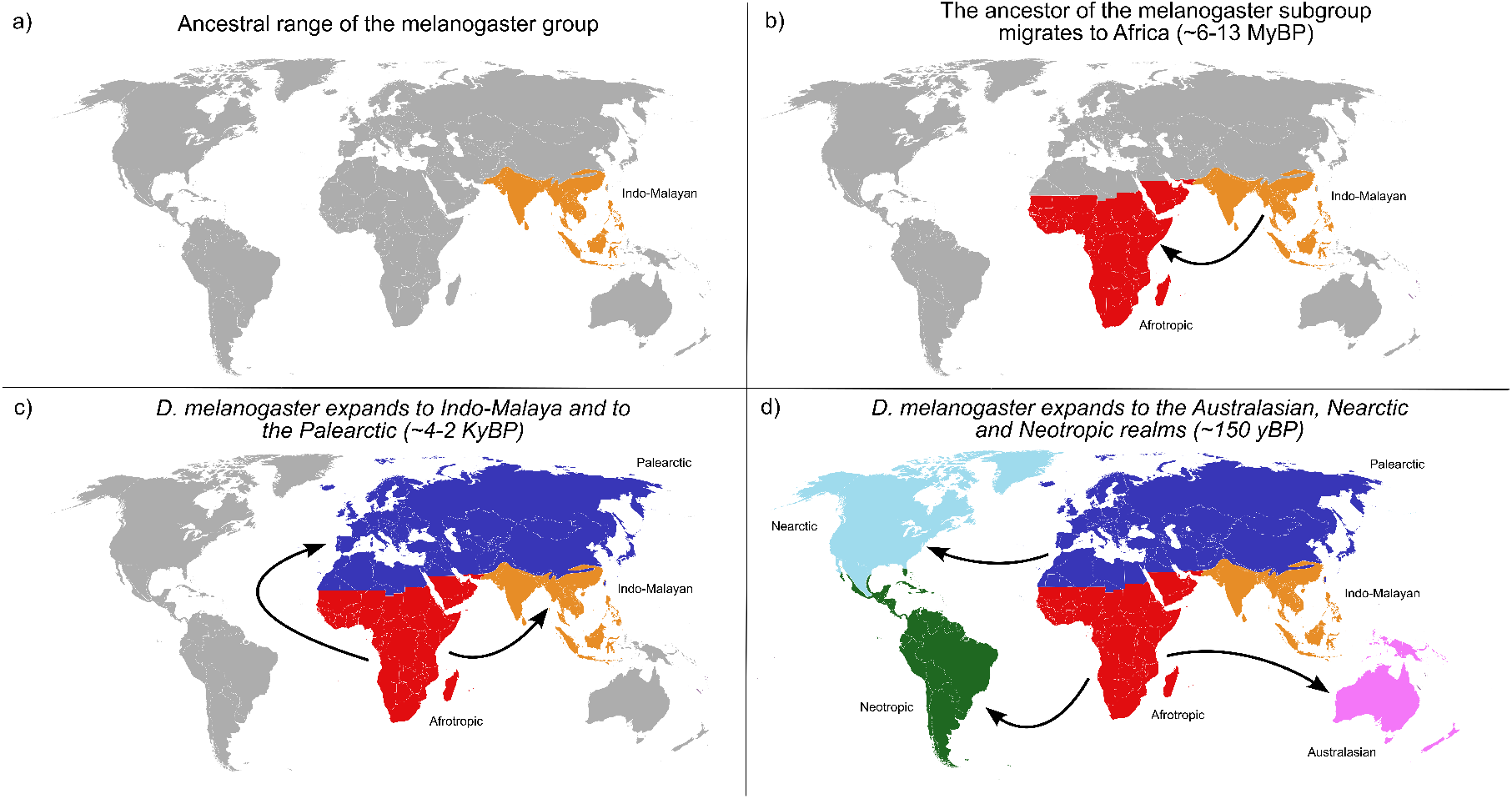
Overview of the biogeographic history of *D. melanogaster*.

The biogeographic and evolutionary history of *D. melanogaster* has been extensively studied; the ancestors of *D. melanogaster* first inhabited part of the Indo-Malayan region [Kopp, 2006], where most of the species of the melanogaster group are found today [Schawaroch, 2002]. At a time roughly estimated between 6-13 Mya, a lineage ancestral to *D. melanogaster* and its closely related species migrated to Sub-Saharan Africa [Russo et al., 1995]. This proto-melanogaster lineage then diverged into 9 species, those that today frame the melanogaster subgroup [David et al., 2007]: *D. orena* and *D. erecta* form the ‘erecta complex’, *D. yakuba, D. santomea* and *D. teissieri* form the ‘yakuba complex’ and *D. mauritiana, D. sechellia, D. simulans* and, eventually, *D. melanogaster*, form the ‘melanogaster complex’ [Ko et al., 2003]. All of these species, with the exception of *D. simulans* and *D. melanogaster* which are now cosmpolitan, are still confined to the Afrotropical region [Markow and O’Grady, 2005]. *D. melanogaster* and its sister species *D. simulans* managed to evolve as human commensals —with different levels of commensalism— and were hence assisted in their spread to the cosmopolitan distributions they exhibit today [Lachaise and Silvain, 2004, Sprengelmeyer et al., 2020, Li and Stephan, 2006]. *D. melanogaster* first expanded its range to the Indo-Malayan realm (∼4000ya [Chen et al., 2024]), followed by the Palearctic (∼2000ya [Chen et al., 2024]) and, only in very recent times [Chen et al., 2024, Arguello et al., 2019, Keller, 2007](∼150-200ya), by the Australasian, Nearctic and Neotropical realms.

In total *D. melanogaster* has about 120-130 TE families Quesneville et al. [2005], Kaminker et al. [2002], 11 of which were recently acquired by HTT (8 from *D. simulans* and 3 from Neotropical species; see above). This raises the question of the origin of the other ≈ 110 TE families that were not subject to HTT during the last 200 years. Previous studies found that most insertions of LTR families in *D. melanogaster* have a very high sequence identity, and are therefore likely to be of very recent origin, possibly as young as 16.000 years old [Bergman and Bensasson, 2007]. Furthermore, insertions of many families segregate at a very low population frequency, suggesting that these families were recently active [Kofler et al., 2012, 2015]. It is possible that recent HTT events triggered the high rate of activity observed for many *D. melanogaster* TEs. Previous studies have identified multiple HTT events between *D. melanogaster, D. simulans* and *D. yakuba* [Bartolomé et al., 2009, Sánchez-Gracia et al., 2005, Loreto et al., 2008]. Another study suggested that several *D. melanogaster* TEs were acquired by HTT from *D. simulans* [Modolo et al., 2014]. *Also D*.*subobscura, D*.*teisseri, D. erecta, D*.*secchelia* and species of the genus *Zaprionus* may have been involved in HTT with *D. melanogaster* [Loreto et al., 2008, Carareto, 2011, Maruyama and Hartl, 1991, de Setta et al., 2011, Simao et al., 2018]. Recently, 285 assemblies of 262 drosophilid species became available [Kim et al., 2021, 2024], allowing us to revisit the question of the origin of *D. melanogaster* TEs. Using this extensive dataset, we identified sequences resembling *D. melanogaster* TEs and inferred likely HTT events, as well as the direction and the biogeographic realm of the horizontal exchange. The TE composition of *D. melanogaster* mirrors its biogeographic history: we find sequences resembling many *D. melanogaster* TEs in Afrotropical species, a few in Indo-Malayan species and for three TEs in Neotropical species. This pattern is likely shaped by HTT. We found a marked asymmetry in the direction of the horizontal exchange: *D. melanogaster* received approximately three times more TEs from other species than it donated to them. We did not find any horizontal exchange of *D. melanogaster* TEs with species from the Nearctic, Palearctic or Australasian regions, even though these regions were colonized by *D. melanogaster* 200-2000 years ago. Extending this analysis to 18 other drosophilids shows that most HTT occurred within the ancestral biogeographic realm of the focal species. We conclude that biogeography is a major factor shaping the TE composition of drosophilds, with a striking impact on their genome evolution.

## 2 Results

### 2.1 The TE composition of *D. melanogaster* mirrors its biogeographic history

Here, our aim is to shed light on the origin of the TE families in *D. melanogaster*. First, we investigated whether other drosophilid species harbour TE insertions resembling those in *D. melanogaster*. To achieve this, we curated a dataset comprising 285 assemblies from 262 drosophilid species [Kim et al., 2021, 2024] (see Supplementary Fig. S2), and inferred the ancestral biogeographic realm for each species [Markow and O’Grady, 2005] (Supplementary File S1). Using RepeatMasker, we identified sequences in these assemblies that were highly similar to *D. melanogaster* TEs. For each TE family, we considered only the best match per assembly, as this is most likely to reflect a HTT event. We quantified similarity between the *D. melanogaster* TE and each candidate using a score based on the Smith-Waterman alignment, which accounts for sequence divergence and alignment length (see Supplementary Fig. S1). A score of *s* = 1.0 indicates an identical match to the *D. melanogaster* consensus, while *s* = 0.0 indicates no similarity at all. Only insertions with high similarity (score ≥ 0.5) were retained for further analysis. Details of the matched sequences, including genomic location, sequence divergence, alignment length, and similarity score, are provided in Supplementary File S2.

We identified numerous sequences that are highly similar to *D. melanogaster* TEs across various drosophilid species (Fig. 2). As expected, closely related species from the melanogaster subgroup exhibit highly similar TE insertions. However, we also found similar insertions in more distantly related species within the melanogaster group, such as *D. vallismaia* and *D. merina*, and even in more distantly related *Zaprionus* species. This is consistent with previous findings of similar TE sequences shared between *Zaprionus* and members of the melanogaster group [Carareto, 2011]. Of particular note is *Scaptodrosophila latifasciaeformis*, the most divergent Afrotropical species from *D. melanogaster*, which harbours insertions similar to 16 *D. melanogaster* TE families.

**Figure 2:**
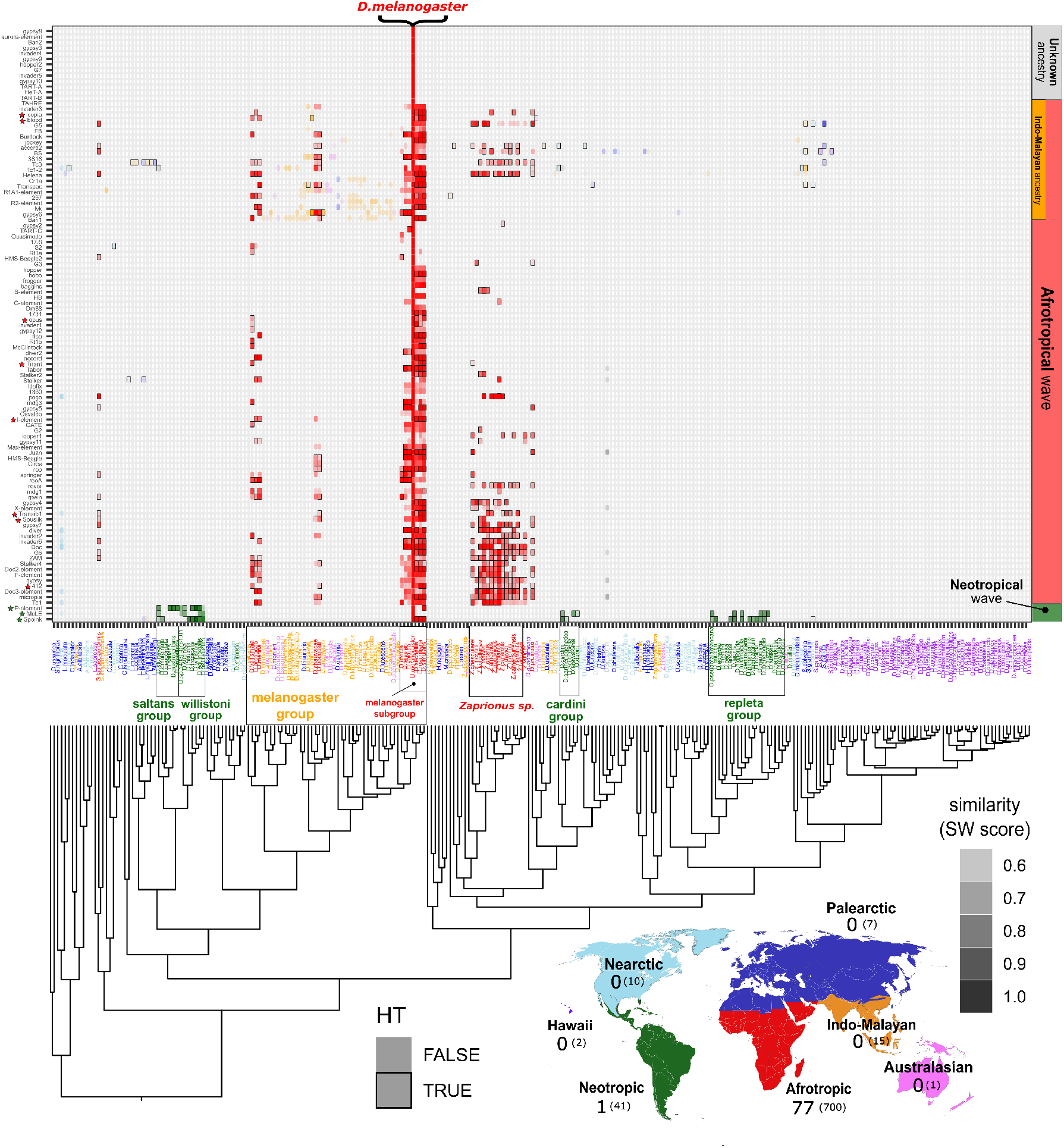
Overview of species that carry sequences resembling *D. melanogaster* TEs. The matrix shows the similarity between *D. melanogaster* TEs (y-axis) and the best matching sequence in each of 262 drosophilids species (x-axis). The color reflects the biogeographic realm and the intensity of the colors scales with the similarity score (see text; solely scores > 0.5 are shown). Black frames indicate likely HTT events with *D. melanogaster* as determined by our formal test. Species were arranged according to their phylogeny and TEs were grouped according to their biogeographic origin (panel at the right). Three distinct groups of TEs are observed: Neotropical (green, 3 families), Afrotropical (red, 92 families) and unknown (grey, 17 families).

When considering biogeographic realms, it is striking that all the investigated drosophilid species from Afrotropical regions contain sequences that closely resemble *D. melanogaster* TEs (Fig. 2). To test whether this pattern extends beyond drosophilids, we examined the genomes of 1369 arthropod species. Outside of drosophilids, we found only a few highly fragmented and divergent insertions for some TEs (e.g., TC3 and Stalker4) (Supplementary Fig. S3). We therefore conclude that sequences closely resembling *D. melanogaster* TEs are found exclusively in drosophilids. For three TE families (*Spoink, McLE, P* -element), we identified highly similar insertions in Neotropical species from the saltans, willistoni, cardini and repleta groups. These TEs have spread across global *D. melanogaster* populations within the last 70 years, likely following horizontal transfer events facilitated by *D. melanogaster* ‘s expansion into the Americas approximately 200 years ago [Daniels et al., 1990, Pianezza et al., 2023, 2024]. Interestingly, we did not detect any insertions resembling *D. melanogaster* TEs in species from the Palearctic, Nearctic, or Hawaiian regions (Fig. 2).

Of the investigated 107 TE families in *D. melanogaster*, similar insertions were found for 92 families in Afrotropical species (Fig. 2). For a subset of 21 of these 92 TE families we also find weakly similar sequences in non-African species of the melanogaster group, primarily from the Indo-Malayan region. Of the 92 Afrotropical TE families, 85 had similar insertions in species of the melanogaster subgroup, 44 in species of the melanogaster group excluding the subgroup, and 39 in species of the Zaprionus genus (Fig. 2). For 14 TEs, no similar insertions were found in any of the surveyed drosophilid species (unknown origin). These include the telomeric TEs (*HeT-A*/*DM06920, TAHRE, TART-A*/*AY561850, TART-B* /*DM14101*), which are known to evolve rapidly, as well as several likely ancient TEs with insertions that are largely fixed in *D. melanogaster* populations (e.g., *gypsy8, hopper2, DMAURA*/*aurora-element, invader4*; [Kofler et al., 2015]). This pattern is consistent across DNA transposons, LTR elements, and non-LTR retrotransposons (Supplementary Fig. S5).

Similar TE sequences in two species may result from HTT or from common descent through vertical inheritance. To identify putative HTTs, we developed a test based on the expectation that TEs involved in HT will show higher sequence identity than orthologous genes [Peccoud et al., 2018, Wallau et al., 2012]. Specifically, we inferred that a TE had been horizontally transferred if its sequence divergence was lower than the 5% quantile of the divergence observed for orthologous genes (Supplementary Fig. S4). This approach is likely to be conservative, as purifying selection is expected to act more strongly on genes than on TEs. In total, we identified 700 instances of TEs in various species that are more similar to *D. melanogaster* TEs than would be expected under vertical inheritance (Fig. 2a; black boxes). The actual number of HTT events required to explain these patterns is likely substantially smaller, since a single HTT event in the common ancestor of certain species could introduce the TE to all their descendants. However, it can conservatively be assumed that at least one HTT event per TE family is necessary to explain this pattern. Thus, we infer that at least one HTT event occurred in 79 TE families. Apart from the TEs with unknown origin, we found no evidence for HTT for *Osvaldo, 1360, Idefix, baggins, Dm88, HB, frogger, hopper, 17*.*6, Quasimodo, R2-element, R1A1-element, Cr1a, invader3, FB*.

In summary, the TE composition of *D. melanogaster* roughly mirrors its biogeographic history. We identified sequences resembling *D. melanogaster* TEs in a few Indo-Malayan species, reflecting the range of the distal ancestors of *D. melanogaster* [Kopp, 2006], *and in many Afrotropical species, which is consistent with the origin of D. melanogaster* in the western Afrotropical regions [Lachaise et al., 1988]. Additionally, we detected highly similar insertions in Neotropical species for three TE families, resulting from horizontal transfers following *D. melanogaster* ‘s expansion into the Americas around 200 years ago. Overall, we identified evidence for HTT in at least 79 TE families. This widespread HTT is likely responsible for the striking similarity in TE composition between *D. melanogaster* and the other Afrotropical drosophilids. Interestingly, we found very few insertions resembling *D. melanogaster* TEs in species from the Nearctic, Palearctic, or Australasian regions, despite *D. melanogaster* having colonized these areas between 2000 and 200 years ago.

### 2.2 Origin of the horizontally transmitted *D. melanogaster* TEs

We suggested that 79 *D. melanogaster* TEs were involved in HTT. To determine the direction of the HTT, we constructed phylogenetic trees for each of these TE families. Sequences resembling the focal TE (minimum insertion length 500 bp; covering at least 50% of the full TE length) were extracted from all species and maximum-likelihood trees were generated using IQ-TREE [Nguyen et al., 2015]. Phylogenetic trees were successfully reconstructed for 76 of the 79 TE families (we could not generate trees for *diver2, R1A1-element* and *gypsy11*).

We inferred the direction of the HTT by manually inspecting each tree. For example, a paraphyletic tree where *D. melanogaster* insertions are nested within sequences from another species, would suggest that the latter species (or a close relative) acted as the donor. Conversely, if sequences from a species are nested within *D. melanogaster* insertions, the species likely received the TE from *D. melanogaster*. For several TEs, the donor could only be narrowed down to a group of closely related species (Fig. 3; indicated by multiple black bars). In other cases, the data suggest that a TE from *D. melanogaster* may have spread to multiple recipient species (Fig. 3; multiple brown bars). The 76 phylogenetic trees, along with inferred donor and recipient species, are provided in Supplementary File S3.

**Figure 3:**
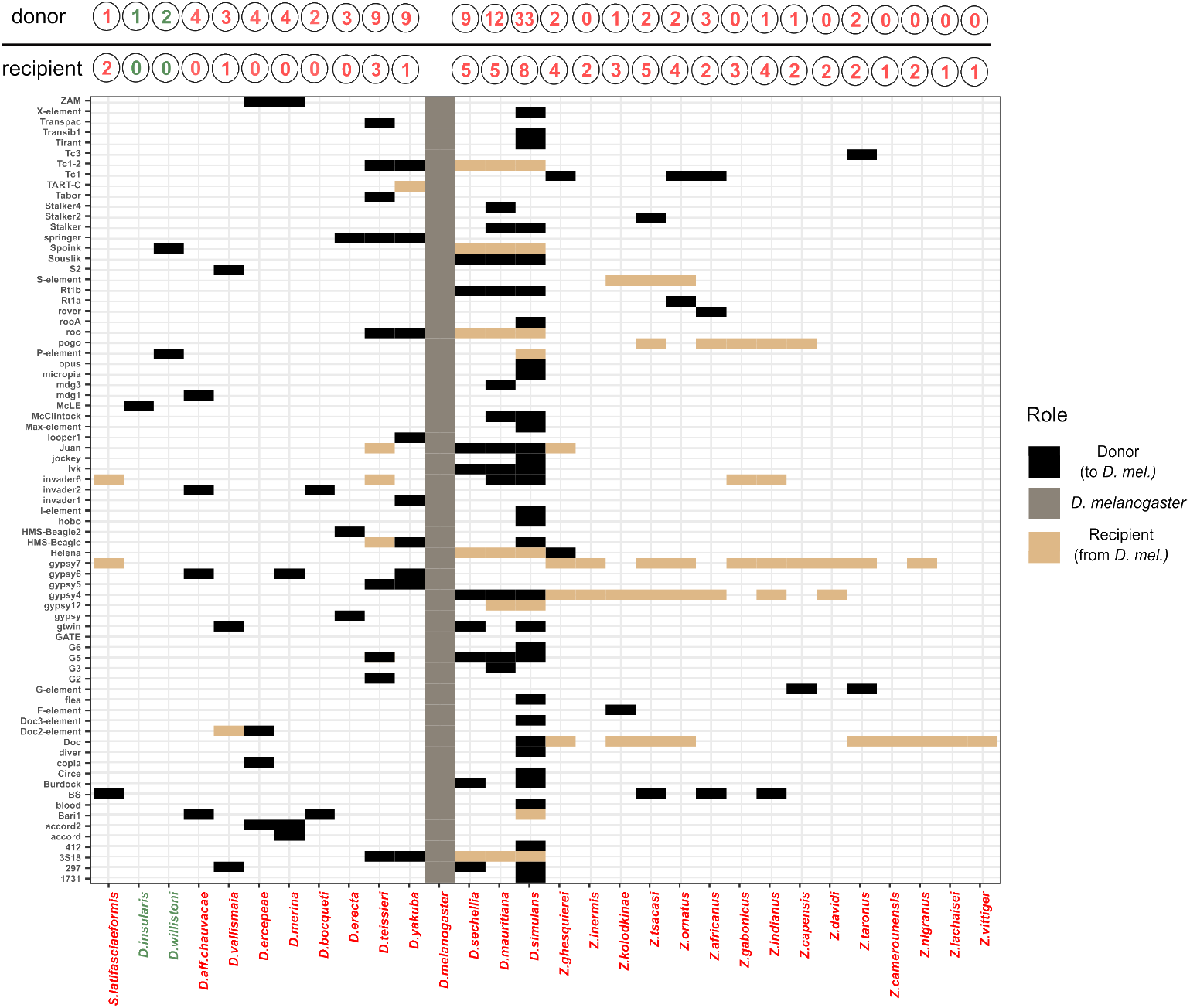
Origin of the horizontally transmitted *D. melanogaster* TEs. Based on phylogenetic trees of the 79 TEs putatively involved in HTT we identified the likely donor of the *D. melanogaster* TEs (black boxes). We also identified the ‘recipient’ in cases where *D. melanogaster* acted as donor (brown boxes). We solely show species involved in at least one HTT. Species are sorted by phylogenetic relatedness (*D. melanogaster* is shown as grey boxes) and colors of the species indicates its biogeographic realm (red: Afrotropical, green: Neotropic).

Based on this approach, we inferred the likely donor species for 70 *D. melanogaster* TEs (Fig. 3; donor). We could not identify the donor for *TARTC-C, Gypsy7, Gypsy12, Pogo, GATE, S-element* (Fig. 3). Our results largely agree with previous studies that inferred the HTT of *D. melanogaster* TEs [Sánchez-Gracia et al., 2005, Bartolomé et al., 2009, Loreto et al., 2008, Modolo et al., 2014, Carareto, 2011]. For a summary of the HTT events identified by previous studies and by our own study, see Supplementary File S4. In several cases, we identified a HTT where previous studies could not. For example, no horizontal exchange of *Tc1* between *D. simulans* and *D. yakuba* could be found [Sánchez-Gracia et al., 2005, Bartolomé et al., 2009]. Consistent with this, we also did not find HTT of *Tc1* between these two species, but our comprehensive analysis of many drosophilid genomes suggests that *Tc1* was horizontally transferred from a species of the genus Zaprionus to *D. melanogaster* (Supplementary File S4). In other cases we reported an alternative donor. For example, previous works suggested a horizontal exchange between *D. simulans* or *D. yakuba* for *Copia* [Sánchez-Gracia et al., 2005, Bartolomé et al., 2009, Modolo et al., 2014] whereas our work suggests that *D. ercepeae* acted as the donor (the insertions between *D. melanogaster* and *D. ercepeae* are 94.67% identical, while only 83.49% identity can be found with *D. melanogaster* and *D. yakuba*). It is not unexpected that our analysis of several hundreds drosophilid genomes would identify alternative donors. However, in some cases, a conflict can also be found, for example, in instances where a previous study found no evidence for a horizontal exchange with *D. simulans* but our work does (Supplementary File S4). This could have several reasons, ranging from biological (the sequence of a TE might differ among strains analysed in different works) to methodological (PCR-based scans might miss insertions found with whole genome analysis, e.g. due to polymorphisms in the primer) to statical (e.g. using different thresholds for inferring HTT).

The closely related *D. simulans* was most frequently involved in HTT with *D. melanogaster*, acting as the donor for 33 TEs. This is consistent with previous studied suggesting that *D. simulans* contributed several TEs to *D. melanogaster* [Scarpa et al., 2023, Pianezza et al., 2024, Schwarz et al., 2021, Carareto, 2011, Blumenstiel, 2019, Bartolomé et al., 2009, Sánchez-Gracia et al., 2005]. We also identified *D. mauritiana* as the donor of 12 TEs, while *D. teissieri, D. yakuba*, and *D. sechellia* each contributed 9 TEs. Note that TE families with multiple possible donors are counted multiple times. Interestingly, our data suggest that the distantly related *D. aff. chauvacae, D. vallismaia*, and *D. ercepeae* each acted as donors for 3–4 TE families. Several species from the genus *Zaprionus* may also have contributed TEs to *D. melanogaster*, consistent with Carareto [2011]. Finally, in line with previous work, we found that Neotropical species were the likely donors of the recent invaders *Spoink*, the *P* -element, and *McLE* [Daniels et al., 1990, Pianezza et al., 2023, 2024].

We also found that *D. melanogaster* likely donated 19 TEs to other drosophilids (Fig. 3). Again, *D. simulans* was most frequently involved, having received 8 TEs from *D. melanogaster*. *Zaprionus* species may also have acquired TEs from *D. melanogaster* (Fig. 3). Interestingly, we observed an asymmetry in the direction of HTT involving *D. melanogaster* and other species. While *D. melanogaster* appears to have acquired at least 70 TEs from other drosophilids, it acted as a donor in only 19 cases. The extent of this asymmetry varies across species. For example, *D. simulans* contributed 33 TEs to *D. melanogaster* but received only 8 TEs in return (Fig. 3). In contrast, species within the genus *Zaprionus* may have received more TEs from *D. melanogaster* than they transmitted back. This pattern is consistent with previous findings that also reported asymmetric HTT involving *D. melanogaster* TEs [Carareto, 2011].

In total, we were able to identify the likely donor species for 70 TEs acquired by *D. melanogaster*. Most of these were transferred from closely related species such as *D. simulans, D. mauritiana*, and *D. teissieri*, although more distantly related taxa, like *D. vallismaia* and members of the Zaprionus group, also contributed multiple TEs. Overall, our results highlight a clear asymmetry in TE exchange: *D. melanogaster* appears to have received roughly three times more TEs via HTT than it donated to other species.

### 2.3 Abundance of HTT among drosophilid species from different biogeographic realms

Consistent with the idea that biogeography plays a major role in shaping the HTT network, the majority of potential HTT events involving *D. melanogaster* TEs occurred with species from the Afrotropics (ancestral range) and the Neotropics (recently colonized). Surprisingly, we found little evidence for HTT with species from the Palearctic, Nearctic, and Australasian regions, despite *D. melanogaster* having established populations there 200 to 2000 years ago. This suggests that certain habitats may be less conducive for HTT among drosophilids. For instance, vectors or environmental conditions suitable for HTT may be confined to a few regions. As another example, the activity of some TEs, such as the *P-element*, increases with temperature [Kofler et al., 2018]. Consequently, the chances of successful HTT may be higher in warmer climates where the TE activity is higher [Le Rouzic and Capy, 2005]. To test this hypothesis, we extended our search for HTT to other drosophilids. We selected three species from each biogeographic realm: Afrotropic (*D. melanogaster, S. lattifasciaeformis, Z. indianus*); East Asia and Oceania (we merged the two realms; *D. ananassae, D. suzukii, D. elegans*); Palearctic (*S. pallida, L. andalusiaca, D. subobscura*); Nearctic (*D. recens, D. virilis, D. robusta*); Neotropic (*D. willistoni, D. saltans, D. repleta*); and Hawaii (*D. picticornis, D. grimshawi, D. fungiperda*). As high-quality repeat libraries are available only for a few model organisms, such as *D. melanogaster*, we *de novo* annotated TEs using EarlGrey [Baril et al., 2024]. For each species, we tested for the presence of sequences resembling the TEs in any of the 262 drosophilid species, and for evidence for HTT.

We first validated this approach using *D. melanogaster*. We tested whether the *de novo* annotation method could reproduce the similarity matrix previously generated using the high-quality annotation of *D. melanogaster* TEs (Fig. 2; [Quesneville et al., 2005]). Using both repeat libraries (EarlGrey and consensus sequences), we observed a very similar pattern (supplementary Fig. S6). Most *D. melanogaster* TEs exhibit insertions with similar sequences in Afrotropical species, several show faint similarities in Indo-Malayan species, and 3–4 families have similar insertions in Neotropical species (Supplementary Fig. S6).

We then inferred TE similarity matrices for all 18 species (see Supplementary File S5). For example, the matrix for *S. pallida*, a cosmopolitan species of Palearctic origin [Zetterstedt, 1842], is shown in Fig. 4a. We found many sequences in Palearctic and Nearctic species with a high similarity to TEs in *S. pallida*, and a smaller number in species from the Neotropics, East Asia, and the Afrotropics (Fig. 4a). Our test for horizontal transfer suggests that 106 out of 356 TEs were likely to be involved in a HTT (Fig. 4a). This is the minimum number of HTT events necessary to explain the observed distribution of sequences resembling TEs in *S. pallida*.

**Figure 4:**
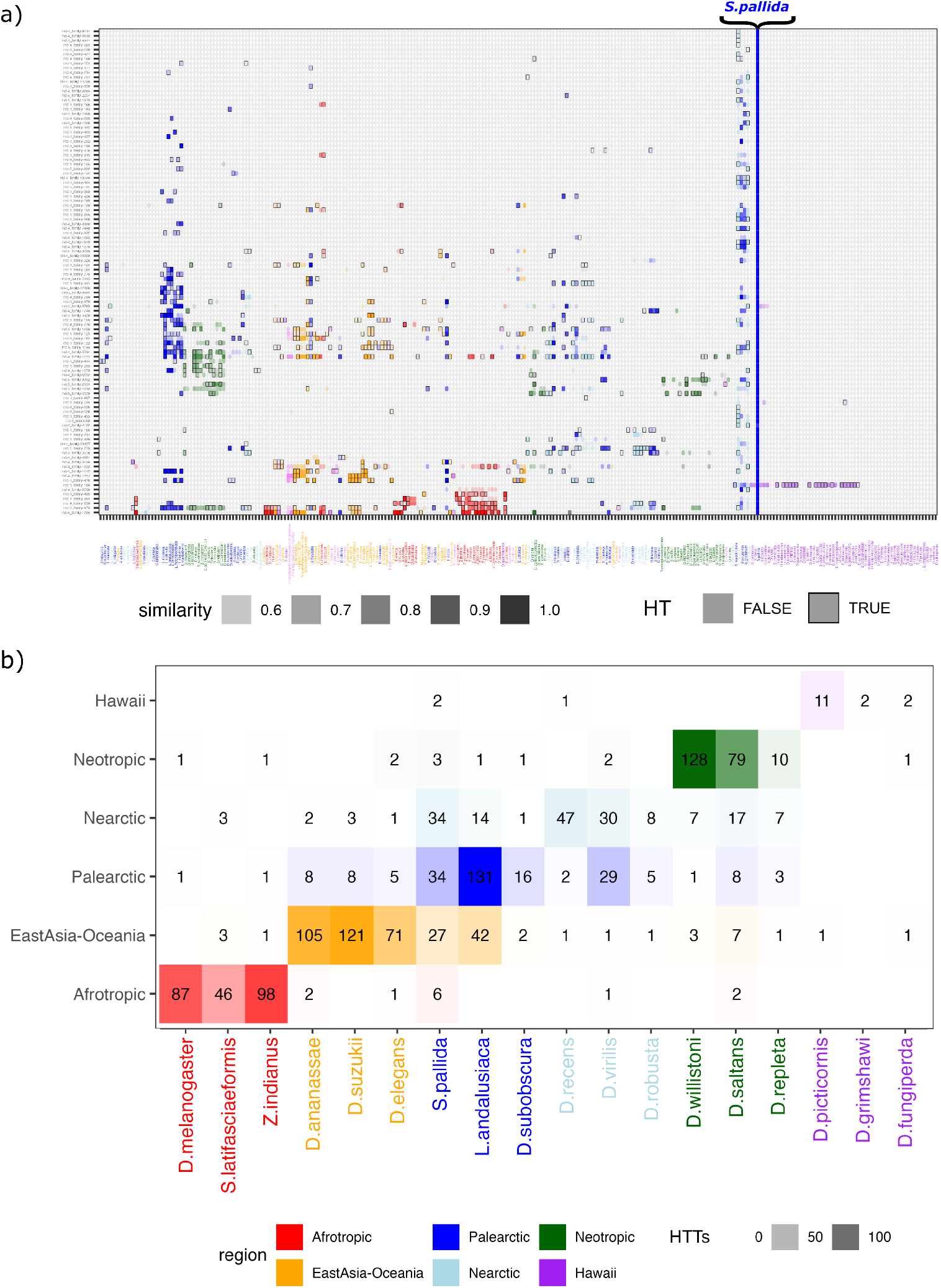
Abundance of HTT among drosophila species from different biogeographic realms. a) Matrix showing the distribution of TE insertions with similarity to *S*.*pallida*. Color intensity scales with similarity and black boxes indicate HT with *S*.*pallida*. Using the best match for each TE we estimate that 34 HTTs occured with Paleartic species, 34 with Nearctic, 27 with Indo-Malaysian b) Summary showing the number of TE families with evidence for HTT in several Drosophila species from each biogeographic realm. HTT event have been classified by biogeographic realm as explained above.

To summarise the biogeographic distribution of HTT events, we assigned each TE to the region where the strongest evidence for HTT was observed (i.e., the highest similarity score). Based on this summary, we estimate that in *S. pallida* many TE families were involved in HTT with Palearctic species (34 families), with Nearctic species (34 families) and species from East Asia (27 families) and a few with Afrotropic species (6 families; Fig. 4a,b).

When this summary is applied to all 18 species, a clear pattern emerges: the majority of HTT events occur within the same biogeographic realm (Fig. 4b). Furthermore, substantial levels of HTT were detected in all regions except Hawaii (Fig. 4b). Some species also show high levels of HTT with species from different realms. For example, *S. pallida* (mentioned above) exhibits multiple HTT events with species from the Palearctic, Nearctic and East Asia. We found that many TEs of Palearctic species are involved in HTT with East Asian species, while few TEs of East-Asian species show HTT with Paleartic species (Fig. 4b). The reason for this asymmetry remains unclear. The rate of HTT between Nearctic and Palearctic species also appears to be high (Fig. 4b). Finally, HTT frequencies also vary within regions. For instance, while *D. willistoni* shows a high number of HTT events within its biogeographic realm (i.e. the Neotropics), *D. repleta* has few (Fig. 4b).

We estimate that the total proportion of nucleotides involved in HTT ranges from 3.3% in *D. fungiperda* to 55% in *D. suzukii*, with a median of 22.1% (Supplementary Table S1). These values are slightly higher than the 2–25% range previously reported for insects [Peccoud et al., 2017].

In summary, we detected substantial levels of HTT in drosophilids across all biogeographic realms except Hawaii. Most HTT events occurred among species within the same biogeographic realm.

### 2.4 Biogeography shapes the TE composition

The presence of similar TE sequences in species from the same biogeographic realm suggests that biogeography plays a major role in shaping the TE landscape. However, this pattern could be confounded by shared ancestry, since closely related species often inhabit the same biogeographic realm and may therefore possess similar TE sequences due to vertical inheritance. To disentangle the effects of shared ancestry from those of shared biogeographic realm, we contrasted the similarity of TEs with that of genes. Since genes are vertically transmitted, their divergence reflects the relatedness of the species. As a proxy, we computed the average divergence of BUSCO genes. To estimate the average TE similarity among species, we used RepeatMasker to identify sequences resembling the TEs of a focal species in all other genomes. We retained the best match for each TE family per assembly.

As expected, in *D. melanogaster* we observed a negative correlation between the gene divergence and the average similarity of TEs (Fig. 5a). Such a correlation is consistent with both vertical transmission and frequent HTT. However, examining the residuals of this correlation revealed that *D. melanogaster* TEs are significantly more similar to TEs in Afrotropical species than to TEs in species from other realms (Fig. 5b; t-test: *p* = 3.5*e* − 12). This elevated TE similarity among Afrotropical species remains significant even when species from the melanogaster subgroup, which are themselves Afrotropical, are excluded (t-test; *p<*2.2*e* − 16).

**Figure 5:**
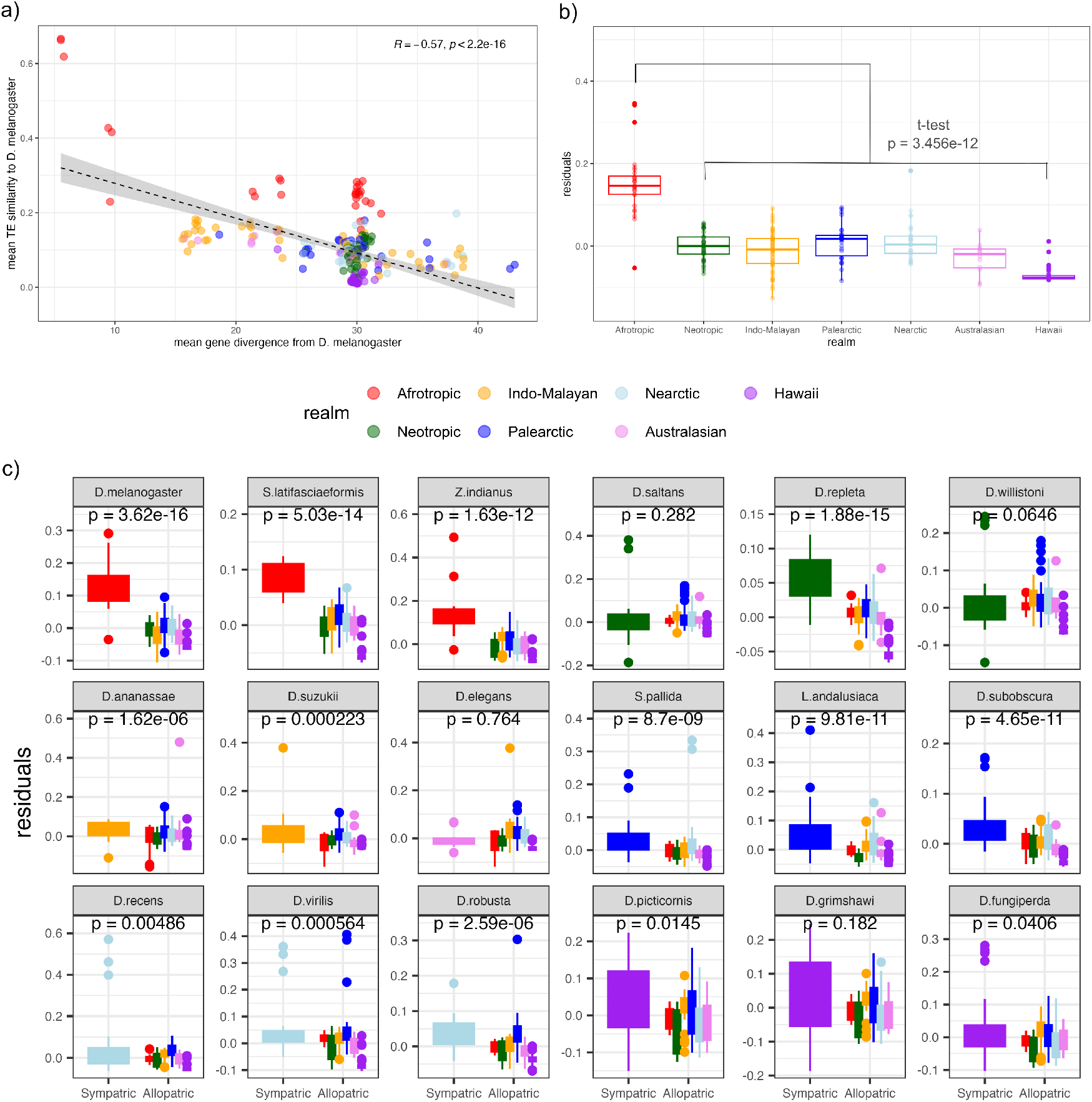
Biogeography is an important factor shaping the TE composition. a) The scatter plot shows the average sequence similarity between *D. melanogaster* TEs (consensus sequences) and TE sequences in 262 drosophilds (y-axis) versus the average divergence of BUSCO genes (x-axis). The average similarity of *D. melanogaster* TEs to insertions in Afrotropical species is elevated compared to other species. b) Residuals of the scatter plot shown in ‘a’ grouped by biogeographic realm. This shows that *D. melanogaster* TEs are on the average more similar to TEs in Afrotropical species than expected based on genetic distance. c) Residuals for 18 drosophilid species from diverse biogeographic realms (TEs annotated with EarlGrey). For 11 out of 18 species the average sequence similarity of the TEs is elevated within species from a biogeographic realm than in species from other realms (*p <* 0.05*/*18; Bonferroni correction).

Next, we extended this analysis to the 18 species from the different biogeographic realms described above (Fig. 5c). For 11 out of the 18 species, TE similarity was significantly higher among species from the same realm than among species from different realms (Figs. 4b, 5c). Species from Hawaii, for which we detected few HTT events, showed the weakest biogeographic signal (figs. 4b, 5c). However, some species with substantial levels of HTT, such as *D. willistoni*, also showed a weak influence of biogeography (figs. 4b, 5c). This may be due to our conservative approach of including species from the same biogeographic realm as the focal species when computing the correlation. For example, in the case of *D. willistoni*, all Neotropical species exhibit a lower genetic divergence than species from other realms, which may bias the correlation and thus lead to the effect of biogeography being underestimated (supplementary Fig. S7).

To control for this potential bias, we repeated the analysis, this time excluding species from the same biogeographic realm as the focal species when computing the correlations. Using this approach, the biogeographic realm significantly impacted the TE similarity of all species, except those from Hawaii (supplementary Fig. S8).

In summary, our results demonstrate that biogeography influences the TE composition of drosophilid genomes, independently of shared ancestry.

## 3 Discussion

The aim of this study was to trace the origin of *D. melanogaster* TEs, and we found strong evidence for substantial HTT between Afrotropical species. Many *D. melanogaster* TEs likely originated from closely related species such as *D. simulans* and *D. mauritiana*, but also from more distantly related taxa, including members of the *Zaprionus* group. Interestingly, we found little evidence for HTT between *D. melanogaster* and species from the Palearctic and Nearctic, despite *D. melanogaster* having colonized these regions approximately 200–2000 years ago. Nevertheless, a broad analysis of 18 drosophilid species from all major biogeographic realms revealed that HTT is frequent in each realm (except for Hawaii). Our findings suggest that biogeography is an important factor in shaping the TE composition of drosophilid genomes.

We investigated patterns of HTT using the genomes of 262 drosophilid species [Kim et al., 2021, 2024]. As the number of available drosophilid genomes is expected to grow in the future, it is important to consider how this might influence our conclusions. Our finding that biogeography is a major force shaping TE composition in drosophilids is unlikely to change, as it is based on broad patterns across many species. However, the inferred donor species for some *D. melanogaster* TEs may shift with the inclusion of additional genomes (Fig. 3). Nevertheless, since donor species are likely to be identified within clades where the TE is ancestral, future updates may be limited to changes of species within the same clade, rather than between distant ones.

Another potential issue is that the inferred biogeography of certain species may be based on incomplete data. Additionally, species change their distribution ranges. For example *D. melanogaster* emerged in the Afrotropics and spread to the rest of the world around 200-2000 years ago. We chose to use the ancestral distribution (e.g. Palearctic for *S. pallida* and Afrotropic for *D. melanogaster*), because the HTT pattern is probably most profoundly influenced by this long-term distribution.

Our work suggests that biogeography shapes the TE composition of drosophilids, raising the question of how an ecological property can influence a genomic feature. Although the precise mechanisms of HTT are unclear, physical proximity between species is likely to enable it, for example via viruses [Reiss et al., 2019, Gilbert et al., 2014]. However, biogeographic overlap alone may not be sufficient, as at least some degree of temporal and ecological overlap is also necessary to enable HTT [Carareto, 2011, Loreto et al., 2008]. This could explain the differences observed in the TE composition between species in the same biogeographic realm. Furthermore, biogeography can only exert a significant impact on the TE composition if the rate of HTT in drosophilids is very high. This is consistent with previous studies that have identified 11 HTT events in the last 200 years [Pianezza et al., 2024, Scarpa et al., 2023] in one species only. In general, it is likely that HTT is very common in insects [Gilbert and Cordaux, 2017].

The finding that most TEs in *D. melanogaster* show signatures of HTT raises the question of whether some TEs are transmitted solely vertically. For example, it is likely that many non-LTR families are largely vertically transmitted [Loreto et al., 2008, Schaack et al., 2010]. However, we found few differences among DNA transposons, LTR and non-LTR retrotransposons in *D. melanogaster* (Supplementary Fig. S5). TE families for which we could not detect evidence for HTT may be predominantly vertically inherited. Nevertheless, it is not clear how vertically inherited TEs could escape gradual erosion of their sequence by mutations. Perhaps a low level of residual activity, despite an active host defence, is capable of preserving a few functional copies of a TE in a species. Whether such residual activity is feasible depends on the efficiency of host defence mechanisms. In *D. melanogaster*, the piRNA pathway silences TEs but it is not clear if the repressed TEs may exhibit some residual activity.

We propose that many TEs found in *D. melanogaster* were acquired from closely related species such as *D. simulans* and *D. mauritiana*. However, this merely shifts the question of origin one step back: where did these species acquire the TEs from? One possibility is that some unassembled or unsampled species acted as the original donors. Alternatively, horizontal transposon transfer (HTT) may frequently occur among closely related species through a “merry-go-round” model, in which the same TE lineage circulates among species over evolutionary time. For example, a TE might first be transferred to *D. melanogaster*, then to *D. mauritiana*, and later reintroduced into *D. melanogaster*. Such a pattern has been proposed for several *D. melanogaster* TEs, including *hobo* and the *I-element* [Blumenstiel, 2019].

Consistent with this idea, many TEs in *D. melanogaster* appear to have spread through multiple invasion waves, each likely triggered by a distinct HTT event [Schwarz et al., 2021, Pianezza et al., 2023, 2024]. The coexistence of multiple TE subfamilies within a single species is a common observation [Clark et al., 1994, Haring et al., 2000, Pinsker et al., 2001, Robertson and MacLeod, 1993], suggesting that HTT, while frequent, often remains limited to a small group of closely related species.

We speculate that host defense mechanisms constrain the reinvasion rate of TEs. In particular, the piRNA pathway silences TEs with sequence complementarity to existing piRNAs [Brennecke et al., 2007, Gunawardane et al., 2007, Gainetdinov et al., 2023]. TEs that have diverged sufficiently —typically by 10–20%— from the existing piRNA pool may evade repression and establish a novel invasion following HTT [Schwarz et al., 2021, Kotov et al., 2019].

We observed a striking asymmetry in the direction of HTT: more TEs were transferred from *D. simulans* (and related species) to *D. melanogaster* than vice versa. What might account for this imbalance? One previously proposed explanation involves differences in effective population size (*N*_*e*_) between species [Carareto, 2011]. In species with large *N*_*e*_, purifying selection is more efficient, potentially removing newly acquired TEs before they can establish. Conversely, in species with smaller *N*_*e*_, genetic drift may overpower selection, allowing TEs to persist and spread. If *D. melanogaster* has a smaller effective population size than *D. simulans*, this could explain why *D. melanogaster* retains more horizontally acquired TEs [Gonzalez and Petrov, 2012, Carareto, 2011]. Another hypothesis is that species differ in their susceptibility to HTT. Differences in the accessibility of the germline to vectors of HTT, such as viruses, mites, or bacteria, may affect how easily TEs can be introduced into new hosts. It is possible that the germline of *D. melanogaster* is more permissive or accessible to such vectors than that of *D. simulans*. A further possibility is that the cost of infection by HTT-mediating vectors varies across species. For example, Wolbachia, an intracellular bacterium suspected to facilitate HTT [Loreto et al., 2008], has been shown to reduce fertility more severely in *D. simulans* than in *D. melanogaster* [Merçot and Charlat, 2004]. Such differences could reduce the likelihood of vector persistence and TE transfer in *D. simulans*, contributing to the observed asymmetry.

For the vast majority of *D. melanogaster* TEs, we find highly similar sequences in other Afrotropical species, indicating a network of species regularly exchanging TEs by HT. The recent colonization of the Neotropics by *D. melanogaster* (and *D. simulans*) has opened a new chapter in colonization of the genomes of these Afrotropical species by TEs. Over the past 200 years, three TEs from Neotropical species have successfully invaded *D. melanogaster*. These TEs are, so far, only detected in these Neotropical species, *D. melanogaster*, and a few other species of the simulans complex (Fig. 2). Given the extensive network of HTT among Afrotropical species it is likely that the Neotropical TEs will gradually spread into other Afrotropical drosophilids. Furthermore, as *D. melanogaster* continues to coexist with Neotropical species, it will likely acquire additional TEs from Neotropical species, that may again spread into Afrotropical species, similarly to the two LTR transposons *Spoink* and *Shellder* [Scarpa et al., 2025]). Vice versa, it is likely that Afrotropical TEs present in genomes of *D. melanogaster* and *D. simulans* will gradually spread into some Neotropical species. Indeed, it was previously suggested that the TE *Copia* was transmitted from *D. melanogaster* to *D. willistoni* [Jordan et al., 1999], although this finding was not confirmed by a subsequent study [Rubin et al., 2011]. We also did not detect a single *Copia* insertion in any of the two analysed *D. willistoni* genome assemblies. In any case it is likely that the habitat expansion of *D. melanogaster* and *D. simulans* connected the previously separated networks of HTT in Afrotopical and Neotropical species. As a result many Afrotropical and Neotropical species may eventually end up with similar sets of TEs.

A puzzling observation is the apparent absence of any recent TE exchange between *D. melanogaster* and species from outside the Afrotropics and Neotropics. Notably, the Indo-Malayan and Palearctic regions have been colonized by *D. melanogaster* for 2000 to 4000 years, while the Nearctic and Australasian regions were colonized more recently (around 200 years ago), roughly at the same time as the colonization of the Neotropics. However, our investigation shows that HTT does occur in drosophilids from these regions. We therefore expect that *D. melanogaster* will eventually also acquire TEs from drosophilids of the Indo-Malayan, Palearctic, Nearctic, and Australasian realms, and vice versa.

In summary, we propose that the global expansion of *D. melanogaster* has initiated a new era of widespread HTT between drosophilid species in contact with *D. melanogaster*. This wave of horizontal exchange may continue until all species connected by a network of HTT share a broadly similar set of TE families.

## 4 Materials and Methods

### 4.1 TE library and assemblies

We obtained a database of the consensus sequences of *D. melanogaster* TEs [Quesneville et al., 2005] and added TEs that recently invaded *D. melanogaster* : *Spoink, McLE, Souslik, Transib1 new* [Pianezza et al., 2024, Ellison and Cao, 2020, Pianezza et al., 2023]. We removed TEs with sequences shorter than 500 bp.

We obtained 279 highly contiguous assemblies of 262 drosophilids species [Kim et al., 2021, 2024] (repository accessed on 01/05/2024) and added 4 *D. melanogaster* assemblies (GCA_003401735.1, GCA_020141585.1, GCA 020141505.1, GCA 020141515.1) and two *D. simulans* assemblies (GCA_039725605.1, GCA_039725755.1 [Signor et al., 2023]).

For each of the 262 species, we manually assigned its ancestral biogeographic realm based on Markow and O’Grady [2005] and if no information was available for a given species, the portal GBIF (https://www.gbif.org/ [GBIF, 2024]).

### 4.2 TE presence/absence and SW similarity score

To screen for the presence of the TEs in the 262 species, we used RepeatMasker (v4.1.2-p1) [Smit et al., 2013-2015] with options ‘-no_is’, ‘-s’ ‘-u’, ‘-nolow’, ‘-lib’, providing our custom library as query. We used the ‘.ori.out’ RM output files to compare the Smith-Waterman (SW) scores of each TE among the species. For each TE, we identified the highest SW score in each assembly and the highest SW score among all the assemblies. For each species we calculated the similarity score as the ratio between the highest score in the given species and the highest score among all species. For the species where several assemblies were available we used the maximum score among the assemblies. We filtered TEs with a score *<* 0.8 in *D. melanogaster* (i.e. TEs not present in *D. melanogaster*).

### 4.3 Phylogenetic tree of 262 drosophilids species

To generate a phylogenetic tree among the 262 drosophilid species we used BUSCO (v5.5.0) [Manni et al., 2021] to extract the sequences of orthologous genes the assemblies. *Anastrepha ludens* was included as outgroup. For species with multiple assemblies, we randomly selected one. To generate an alignment of all the shared orthologous genes, the BUSCO output folder was directly used as input for the BUSCO phylogenomics tool [McGowan, 2023] with option ‘–supermatrix_only’ and ‘-psc 97’. The final bayesian tree was obtained by providing the alignment of the genes to BEAST (v2.7.5) [Bouckaert et al., 2019].

### 4.4 Phylogenetic trees of TE insertions

For each TE family, we extracted the sequences of TE insertions in each of the 285 assemblies with bedtools (v2.30.0) [Quinlan and Hall, 2010] (‘getfasta’ function, ‘-s’ option), based on the coordinates found by RepeatMasker. We filtered for insertions with length >0.5 and sequence identity >0.5 to the consensus. We only used one assembly for species with multiple assemblies. We performed a multiple sequence alignment of the TE insertions with using MAFFT (v7.526) [Katoh et al., 2002]. The trees were generated with IQtree (v1.6.12) [Nguyen et al., 2015] using the options ‘-m MFP’, ‘-bb 1000’, ‘-alrt 1000’. We annotated the trees using ape (v5.7-1) [Paradis et al., 2004].

### 4.5 Test for HTT

For each BUSCO gene, we computed the sequence divergence among all pairs of species. We aligned pairs of genes using MAFFT, and calculated the sequence divergence with a custom Python script (genes-divergence.py). We then compared the sequence divergence of the TEs (based on the best match between two species identified by RepeatMasker) to the divergence of the BUSCO genes. We assumed HTT if the divergence of the TE was less than for 95% of the BUSCO genes.

### 4.6 TE annotation

To generate TE libraries for 18 drosophilid species we used EarlGrey (v4.4.5) [Baril et al., 2024] with option ‘-c yes’. Sequences classified as ‘Unknown’, ‘Simple Repeats’, ‘rRNA’ and ‘Satellites’ were excluded from the analysis.

## Acknowledgments

R.P. would like to thank Matthew P. Beaumont and Almorò Scarpa for the fruitful discussions that improved the quality of this work, together with the members of the Institute of Population Genetics.

## Author contributions

R.P. conceived the project and analysed the data. R.P. and R.K. wrote the manuscript. R.K. supervised the project.

## Data Availability

The analysis performed in this work have been documented with RMarkdown and have been made publicly available, together with the resulting figures, at GitHub (https://github.com/rpianezza/Drosophilids-TE-biogeography The final library of *D. melanogaster* TE families consensus sequences used in this work, together with the 18 libraries generated with EarlGrey are also available.

## References

Thomas Wicker, François Sabot, Aurélie Hua-Van, Jeffrey L Bennetzen, Pierre Capy, Boulos Chalhoub, Andrew Flavell, Philippe Leroy, Michele Morgante, Olivier Panaud, et al. A unified classification system for eukaryotic transposable elements. Nature Reviews Genetics, 8(12): 973–982, 2007.

David J Finnegan. Eukaryotic transposable elements and genome evolution. Trends in Genetics, 5(4): 103–107, 1989.

Samuel Aparicio, Jarrod Chapman, Elia Stupka, Nik Putnam, Jer-ming Chia, Paramvir Dehal, Alan Christof-fels, Sam Rash, Shawn Hoon, Arian Smit, et al. Whole-genome shotgun assembly and analysis of the genome of fugu rubripes. Science, 297(5585): 1301–1310, 2002.

Michelle C Stitzer, Sarah N Anderson, Nathan M Springer, and Jeffrey Ross-Ibarra. The genomic ecosystem of transposable elements in maize. PLoS Genetics, 17(10):e1009768, 2021.

Sergey Nurk, Sergey Koren, Arang Rhie, Mikko Rautiainen, Andrey V Bzikadze, Alla Mikheenko, Mitchell R Vollger, Nicolas Altemose, Lev Uralsky, Ariel Gershman, et al. The complete sequence of a human genome. Science, 376(6588): 44–53, 2022.

Peter Sarkies, Murray E Selkirk, John T Jones, Vivian Blok, Thomas Boothby, Bob Goldstein, Ben Hanelt, Alex Ardila-Garcia, Naomi M Fast, Phillip M Schiffer, Christopher Kraus, Mark J Taylor, Georgios Koutsovoulos, Mark L Blaxter, and Eric A Miska. Ancient and novel small RNA pathways compensate for the loss of piRNAs in multiple independent nematode lineages. PLoS Biol., 13(2): 1–20, 2015.

Julius Brennecke, Alexei A Aravin, Alexander Stark, Monica Dus, Manolis Kellis, Ravi Sachidanandam, and Gregory J Hannon. Discrete small RNA-generating loci as master regulators of transposon activity in Drosophila. Cell, 128(6): 1089–1103, 2007.

Benjamin Czech and Gregory J Hannon. One loop to rule them all: the ping-pong cycle and pirna-guided silencing. Trends in biochemical sciences, 41(4): 324–337, 2016.

Justin P Blumenstiel. Birth, school, work, death, and resurrection: the life stages and dynamics of transposable element proliferation. Genes, 10(5): 336, 2019.

Sarah Schaack, Clément Gilbert, and Cédric Feschotte. Promiscuous dna: horizontal transfer of transposable elements and why it matters for eukaryotic evolution. Trends in ecology & evolution, 25(9): 537–546, 2010.

Hua-Hao Zhang, Jean Peccoud, Min-Rui-Xuan Xu, Xiao-Gu Zhang, and Clément Gilbert. Horizontal transfer and evolution of transposable elements in vertebrates. Nature communications, 11(1): 1362, 2020.

ELS Loreto, CMA Carareto, and et P Capy. Revisiting horizontal transfer of transposable elements in drosophila. Heredity, 100(6): 545–554, 2008.

Patrick J Keeling and Jeffrey D Palmer. Horizontal gene transfer in eukaryotic evolution. Nature Reviews Genetics, 9(8): 605–618, 2008.

Jean Peccoud, Vincent Loiseau, Cordaux, and Clément Gilbert. Massive horizontal transfer of transposable elements in insects. Proc Natl Acad Sci U S A, 114(18): 4721–26, 2017.

Almorò Scarpa, Riccardo Pianezza, Filip Wierzbicki, and Robert Kofler. Genomes of historical specimens reveal multiple invasions of ltr retrotransposons in drosophila melanogaster populations during the 19th century. bioRxiv, 2023. doi: 10.1101/2023.06.06.543830.

Florian Schwarz, Filip Wierzbicki, Kirsten-André Senti, and Robert Kofler. Tirant Stealthily Invaded Natural Drosophila melanogaster Populations during the Last Century. Molecular Biology and Evolution, 38(4): 1482–1497, 2021.

A Bucheton, C Vaury, M C Chaboissier, P Abad, A Pélisson, and M Simonelig. I elements and the Drosophila genome. Genetica, 86(1–3): 175–90, 1992.

Georges Periquet, Marie H Hamelin, Yves Bigot, and Antoine Lepissier. Geographical and historical patterns of distribution of hobo elements in Drosophila melanogaster populations. Journal of Evolutionary Biology, 2(3): 223–229, 1989.

Eric Bonnivard, Claude Bazin, Beatrice Denis, and Dominique Higuet. A scenario for the hobo transposable element invasion, deduced from the structure of natural populations of Drosophila melanogaster using tandem TPE repeats. Genetical Research, 75(1): 13–23, 2000.

M. G. Kidwell. Evolution of hybrid dysgenesis determinants in Drosophila melanogaster. Proceedings of the National Academy of Sciences, 80(6): 1655–1659, 1983.

D Anxolabéhère, M G Kidwell, and G Periquet. Molecular characteristics of diverse populations are consistent with the hypothesis of a recent invasion of Drosophila melanogaster by mobile P elements. Molecular Biology and Evolution, 5(3): 252–269, 1988.

S B Daniels, K R Peterson, L D Strausbaugh, M G Kidwell, and A Chovnick. Evidence for horizontal transmission of the p transposable element between drosophila species. Genetics, 124(2):339–355, February 1990. ISSN 1943-2631. doi: 10.1093/genetics/124.2.339. URL http://dx.doi.org/10.1093/genetics/124.2.339.

Riccardo Pianezza, Almorò Scarpa, Prakash Narayanan, Sarah Signor, and Robert Kofler. Spoink, a ltr retrotransposon, invadedd. melanogasterpopulations in the 1990s. bioRxiv, November 2023. doi: 10.1101/2023.10.30.564725. URL http://dx.doi.org/10.1101/2023.10.30.564725.

Riccardo Pianezza, Almorò Scarpa, Anna Haider, Sarah Signor, and Robert Kofler. Unveiling the complete invasion history of d. melanogaster: three horizontal transfers of transposable elements in the last 30 years. bioRxiv, 2024. URL https://www.biorxiv.org/content/early/2024/04/28/2024.04.25.591091.

Pierre Capy and Patricia Gibert. Drosophila melanogaster, drosophila simulans: so similar yet so different. Drosophila melanogaster, Drosophila simulans: So Similar, So Different, pages 5–16, 2004.

Junhao Chen, Chenlu Liu, Weixuan Li, Wenxia Zhang, Yirong Wang, Andrew G Clark, and Jian Lu. From sub-saharan africa to china: Evolutionary history and adaptation of drosophila melanogaster revealed by population genomics. Science Advances, 10(16):eadh3425, 2024.

Artyom Kopp. Basal relationships in the drosophila melanogaster species group. Molecular Phylogenetics and Evolution, 39(3): 787–798, 2006.

Valerie Schawaroch. Phylogeny of a paradigm lineage: the drosophila melanogaster species group (diptera: Drosophilidae). Biological Journal of the Linnean Society, 76(1): 21–37, 2002.

CA Russo, Naoko Takezaki, and Masatoshi Nei. Molecular phylogeny and divergence times of drosophilid species. Molecular biology and evolution, 12(3): 391–404, 1995.

Jean R David, Françoise Lemeunier, Leonidas Tsacas, and Amir Yassin. The historical discovery of the nine species in the drosophila melanogaster species subgroup. Genetics, 177(4): 1969–1973, 2007.

Wen-Ya Ko, Ryan M David, and Hiroshi Akashi. Molecular phylogeny of the drosophila melanogaster species subgroup. Journal of Molecular Evolution, 57: 562–573, 2003.

Therese A Markow and Patrick O’Grady. Drosophila: A guide to species identification and use. 2005.

Daniel Lachaise and Jean-François Silvain. How two afrotropical endemics made two cosmopolitan human commensals: the drosophila melanogaster–d. simulans palaeogeographic riddle. Genetica, 120: 17–39, 2004.

Quentin D Sprengelmeyer, Suzan Mansourian, Jeremy D Lange, Daniel R Matute, Brandon S Cooper, Erling V Jirle, Marcus C Stensmyr, and John E Pool. Recurrent collection of drosophila melanogaster from wild african environments and genomic insights into species history. Molecular Biology and Evolution, 37 (3):627–638, 2020.

Haipeng Li and Wolfgang Stephan. Inferring the demographic history and rate of adaptive substitution in drosophila. PLoS genetics, 2(10):e166, 2006.

J Roman Arguello, Stefan Laurent, and Andrew G Clark. Demographic history of the human commensal drosophila melanogaster. Genome biology and evolution, 11(3): 844–854, 2019.

Andreas Keller. Drosophila melanogaster’s history as a human commensal. Current biology, 17(3):R77–R81, 2007.

Hadi Quesneville, Casey M Bergman, Olivier Andrieu, Delphine Autard, Danielle Nouaud, Michael Ashburner, and Dominique Anxolabehere. Combined evidence annotation of transposable elements in genome sequences. PLoS Comp. Biol., 1(2): 166–175, 2005.

Joshua S Kaminker, Casey M Bergman, Brent Kronmiller, Joseph Carlson, Robert Svirskas, Sandeep Patel, Erwin Frise, David A Wheeler, Suzanna E Lewis, Gerald M Rubin, et al. The transposable elements of the drosophila melanogaster euchromatin: a genomics perspective. Genome biology, 3: 1–20, 2002.

Casey M Bergman and Douda Bensasson. Recent ltr retrotransposon insertion contrasts with waves of non-ltr insertion since speciation in drosophila melanogaster. Proceedings of the National Academy of Sciences, 104(27): 11340–11345, 2007.

Robert Kofler, Andrea J Betancourt, and Christian Schlötterer. Sequencing of pooled dna samples (pool-seq) uncovers complex dynamics of transposable element insertions in drosophila melanogaster. PLoS genetics, 8(1):e1002487, 2012.

Robert Kofler, Tom Hill, Viola Nolte, Andrea Betancourt, and Christian Schlötterer. The recent invasion of natural Drosophila simulans populations by the P-element. PNAS, 112(21): 6659–6663, 2015.

Carolina Bartolomé, Xabier Bello, and Xulio Maside. Widespread evidence for horizontal transfer of transposable elements across drosophila genomes. Genome biology, 10: 1–11, 2009.

Alejandro Sánchez-Gracia, Xulio Maside, and Brian Charlesworth. High rate of horizontal transfer of transposable elements in drosophila. Trends in genetics, 21(4): 200–203, 2005.

Laurent Modolo, Franck Picard, and Emmanuelle Lerat. A new genome-wide method to track horizontally transferred sequences: application to drosophila. Genome biology and evolution, 6(2): 416–432, 2014.

Claudia MA Carareto. Tropical africa as a cradle for horizontal transfers of transposable elements between species of the genera drosophila and zaprionus. Mobile Genetic Elements, 1(3): 179–186, 2011.

Kyoko Maruyama and Daniel L Hartl. Evidence for interspecific transfer of the transposable element mariner between drosophila and zaprionus. Journal of molecular evolution, 33: 514–524, 1991.

Nathalia de Setta, Marie-Anne Van Sluys, Pierre Capy, and Claudia Marcia Aparecida Carareto. Copia retrotransposon in the zaprionus genus: another case of transposable element sharing with the drosophila melanogaster subgroup. Journal of molecular evolution, 72: 326–338, 2011.

Maryanna C Simao, Annabelle Haudry, Adriana Granzotto, Nathalia De Setta, and Claudia MA Carareto. Helena and bs: two travellers between the genera drosophila and zaprionus. Genome Biology and Evolution, 10(10): 2671–2685, 2018.

Bernard Y Kim, Jeremy R Wang, Danny E Miller, Olga Barmina, Emily Delaney, Ammon Thompson, Aaron A Comeault, David Peede, Emmanuel RR D’Agostino, Julianne Pelaez, et al. Highly contiguous assemblies of 101 drosophilid genomes. Elife, 10:e66405, 2021.

Bernard Y Kim, Hannah R Gellert, Samuel H Church, Anton Suvorov, Sean S Anderson, Olga Barmina, Sofia G Beskid, Aaron A Comeault, K Nicole Crown, Sarah E Diamond, et al. Single-fly genome assemblies fill major phylogenomic gaps across the drosophilidae tree of life. PLoS Biology, 22(7):e3002697, 2024.

Jean Peccoud, Richard Cordaux, and Clément Gilbert. Analyzing horizontal transfer of transposable elements on a large scale: Challenges and prospects. Bioessays, 40(2), 2018.

Gabriel Luz Wallau, Mauro Freitas Ortiz, and Elgion Lucio Silva Loreto. Horizontal transposon transfer in eukarya: detection, bias, and perspectives. Genome Biol. Evol., 4(8):689–699, July 2012.

Daniel Lachaise, Marie-Louise Cariou, Jean R David, Françoise Lemeunier, Léonidas Tsacas, and Michael Ashburner. Historical biogeography of the drosophila melanogaster species subgroup. Evolutionary biology, pages 159–225, 1988.

Lam-Tung Nguyen, Heiko A Schmidt, Arndt Von Haeseler, and Bui Quang Minh. Iq-tree: a fast and effective stochastic algorithm for estimating maximum-likelihood phylogenies. Molecular biology and evolution, 32 (1):268–274, 2015.

Robert Kofler, Kirsten-Andre Senti, Viola Nolte, Ray Tobler, and Christian Schlötterer. Molecular dissection of a natural transposable element invasion. Genome research, 28(6): 824–835, 2018.

Arnaud Le Rouzic and Pierre Capy. The first steps of transposable elements invasion: parasitic strategy vs. genetic drift. Genetics, 169(2): 1033–1043, 2005.

Tobias Baril, James Galbraith, and Alex Hayward. Earl grey: a fully automated user-friendly transposable element annotation and analysis pipeline. Molecular Biology and Evolution, 41(4):msae068, 2024.

Johan Wilhelm Zetterstedt. Diptera Scandinaviæ disposita et descripta. Auctore Ph. D: re Johanne Wilhelmo Zetterstedt. ex Officina Lundbergiana, sumtibus auctoris,, 1842. doi: 10.5962/bhl.title.8143. URL http://dx.doi.org/10.5962/bhl.title.8143.

Daphné Reiss, Gladys Mialdea, Vincent Miele, Damien M de Vienne, Jean Peccoud, Clément Gilbert, Laurent Duret, and Sylvain Charlat. Global survey of mobile dna horizontal transfer in arthropods reveals lepidoptera as a prime hotspot. PLoS genetics, 15(2):e1007965, 2019.

Clément Gilbert, Aurélien Chateigner, Lise Ernenwein, Valérie Barbe, Annie Bézier, Elisabeth A Herniou, and Richard Cordaux. Population genomics supports baculoviruses as vectors of horizontal transfer of insect transposons. Nature communications, 5(1): 3348, 2014.

Clément Gilbert and Richard Cordaux. Viruses as vectors of horizontal transfer of genetic material in eukaryotes. Current Opinion in Virology, 25:16–22, August 2017. ISSN 1879-6257. doi: 10.1016/j.coviro.2017.06.005. URL http://dx.doi.org/10.1016/j.coviro.2017.06.005.

Jonathan B Clark, Wayne P Maddison, and Margaret G Kidwell. Phylogenetic analysis supports horizontal transfer of p transposable elements. Molecular biology and evolution, 11(1): 40–50, 1994.

Elisabeth Haring, Sylvia Hagemann, and Wilhelm Pinsker. Ancient and recent horizontal invasions of drosophilids by p elements. Journal of molecular evolution, 51: 577–586, 2000.

Wilhelm Pinsker, Elisabeth Haring, Sylvia Hagemann, and Wolfgang J Miller. The evolutionary life history of p transposons: from horizontal invaders to domesticated neogenes. Chromosoma, 110: 148–158, 2001.

HM Robertson and EG MacLeod. Five major subfamilies of mariner transposable elements in insects, including the mediterranean fruit fly, and related arthropods. Insect Molecular Biology, 2(3): 125–139, 1993.

Lalith S Gunawardane, Kuniaki Saito, Kazumichi M Nishida, Keita Miyoshi, Yoshinori Kawamura, Tomoko Nagami, Haruhiko Siomi, and Mikiko C Siomi. A slicer-mediated mechanism for repeat-associated sirna 5’end formation in drosophila. science, 315(5818): 1587–1590, 2007.

Ildar Gainetdinov, Joel Vega-Badillo, Katharine Cecchini, Ayca Bagci, Cansu Colpan, Dipayan De, Shannon Bailey, Amena Arif, Pei-Hsuan Wu, Ian J MacRae, et al. Relaxed targeting rules help piwi proteins silence transposons. Nature, 619(7969): 394–402, 2023.

Alexei A Kotov, Vladimir E Adashev, Baira K Godneeva, Maria Ninova, Aleksei S Shatskikh, Sergei S Bazylev, Alexei A Aravin, and Ludmila V Olenina. pirna silencing contributes to interspecies hybrid sterility and reproductive isolation in drosophila melanogaster. Nucleic acids research, 47(8): 4255–4271, 2019.

Josefa Gonzalez and Dmitri A Petrov. Evolution of genome content: population dynamics of transposable elements in flies and humans. Evolutionary genomics: statistical and computational methods, volume 1, pages 361–383, 2012.

Hervé Mercot and Sylvain Charlat. Wolbachia infections in drosophila melanogaster and d. simulans: polymorphism and levels of cytoplasmic incompatibility. Drosophila melanogaster, Drosophila simulans: So Similar, So Different, pages 51–59, 2004.

Almorò Scarpa, Riccardo Pianezza, Hannah R Gellert, Anna Haider, Bernard Y Kim, Eric C Lai, Robert Kofler, and Sarah Signor. Double trouble: two retrotransposons triggered a cascade of invasions in drosophila species within the last 50 years. Nature Communications, 16(1): 516, 2025.

I King Jordan, Lilya V Matyunina, and John F McDonald. Evidence for the recent horizontal transfer of long terminal repeat retrotransposon. Proceedings of the National Academy of Sciences, 96(22): 12621–12625, 1999.

PM Rubin, ELS Loreto, CMA Carareto, and VLS Valente. The copia retrotransposon and horizontal transfer in drosophila willistoni. Genetics Research, 93(3): 175–180, 2011.

Christopher E Ellison and Weihuan Cao. Nanopore sequencing and Hi-C scaffolding provide insight into the evolutionary dynamics of transposable elements and piRNA production in wild strains of Drosophila melanogaster. Nucleic Acids Research, 48(1): 1–14, 2020.

S Signor, J Vedanayagam, BY Kim, F Wierzbicki, R Kofler, and EC Lai. Rapid evolutionary diversification of the flamenco locus across simulans clade drosophila species. PLoS Genet, 19, 2023.

GBIF. Gbif home page, 2024. URL https://www.gbif.org. Accessed: 01-05-2024.

A. F. A. Smit, R. Hubley, and P. Green. RepeatMasker Open-4. 0, 2013-2015. URL http://www.repeatmasker.org.

Mosè Manni, Matthew R Berkeley, Mathieu Seppey, and Evgeny M Zdobnov. Busco: assessing genomic data quality and beyond. Current Protocols, 1(12):e323, 2021.

J McGowan. Buscophylogenomics. https://github.com/jamiemcg/BUSCO_phylogenomics, 2023.

Remco Bouckaert, Timothy G. Vaughan, Joëlle Barido-Sottani, Sebastián Duchene, Mathieu Fourment, Alexandra Gavryushkina, Joseph Heled, Graham Jones, Denise Kühnert, Nicola De Maio, Michael Matschiner, Fábio K. Mendes, Nicola F. Müller, Huw A. Ogilvie, Louis du Plessis, Alex Popinga, Andrew Rambaut, David Rasmussen, Igor Siveroni, Marc A. Suchard, Chieh-Hsi Wu, Dong Xie, Chi Zhang, Tanja Stadler, and Alexei J. Drummond. BEAST 2.5: An advanced software platform for bayesian evolutionary analysis. PLOS Computational Biology, 15(4):e1006650, April 2019.

Aaron R Quinlan and Ira M Hall. Bedtools: a flexible suite of utilities for comparing genomic features. Bioinformatics, 26(6): 841–842, 2010.

Kazutaka Katoh, Kazuharu Misawa, Kei-ichi Kuma, and Takashi Miyata. Mafft: a novel method for rapid multiple sequence alignment based on fast fourier transform. Nucleic acids research, 30(14): 3059–3066, 2002.

Emmanuel Paradis, Julien Claude, and Korbinian Strimmer. Ape: analyses of phylogenetics and evolution in r language. Bioinformatics, 20(2): 289–290, 2004.

