## Supplementary Material for "Biogeography shapes the TE landscape of *Drosophila melanogaster*"

### Supplementary figures and tables

#### Supplementary figures

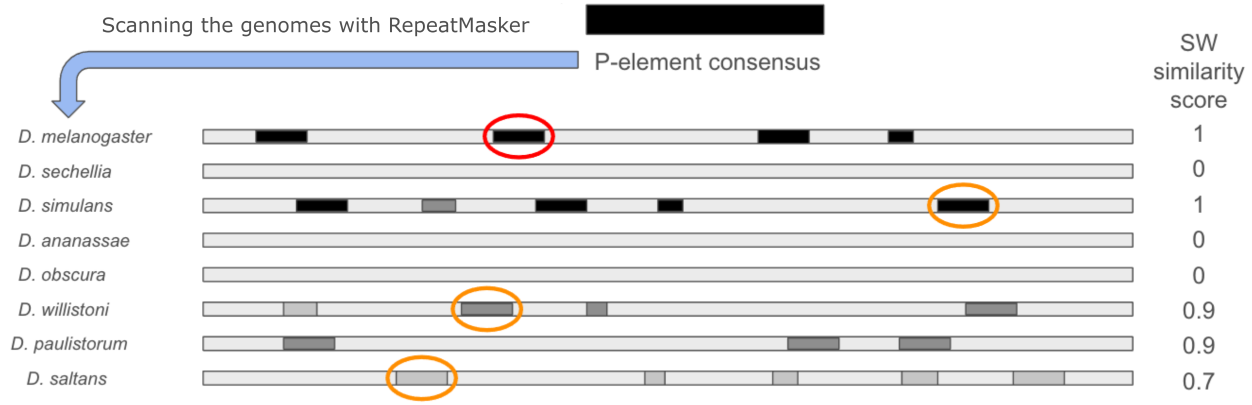


Figure 1: Overview of our approach for finding insertions with a high similarity to *D. melanogaster* TEs in different Drosophilids. The consensus sequences of *D. melanogaster* TEs are used as a query for scanning the genomes of different species with RepeatMasker [Smit et al., 2013-2015]. In each assembly we identify the best matching sequence based on the Smith-Waterman score (colored circle). The SW score reflects both, the length and the sequence similarity of a match. Finally we compute for each TE the similarity score as $score_{TE} = s^{max}_{species}/s^{max}_{all}$ where $s^{max}_{species}$ is the SW score of the best hit in a given species (e.g. orange circle) and $s^{max}_{all}$ the score of the best hit in any species (e.g. red circle, including *D. melanogaster* where typically the best match is expected). We illustrate this approach using the *P*-element as example. The *P*-element in *D. simulans* is identical to the *P*-element in *D. melanogaster* ($score = 1$). We assume, solely as an example to demonstrate the approach, that a slightly different *P*-element sequence is found in *D. willistoni* and *D. paulistorum* ($score = 0.9$), a more diverged sequence one in *D. saltans* ($score = 0.7$) and no matches are found in *D. sechellia*, *D. ananassae* and *D. obscura* ($score = 0.0$). The grey scale of matching regions indicates the similarity to the consensus sequence of the *P*-element.

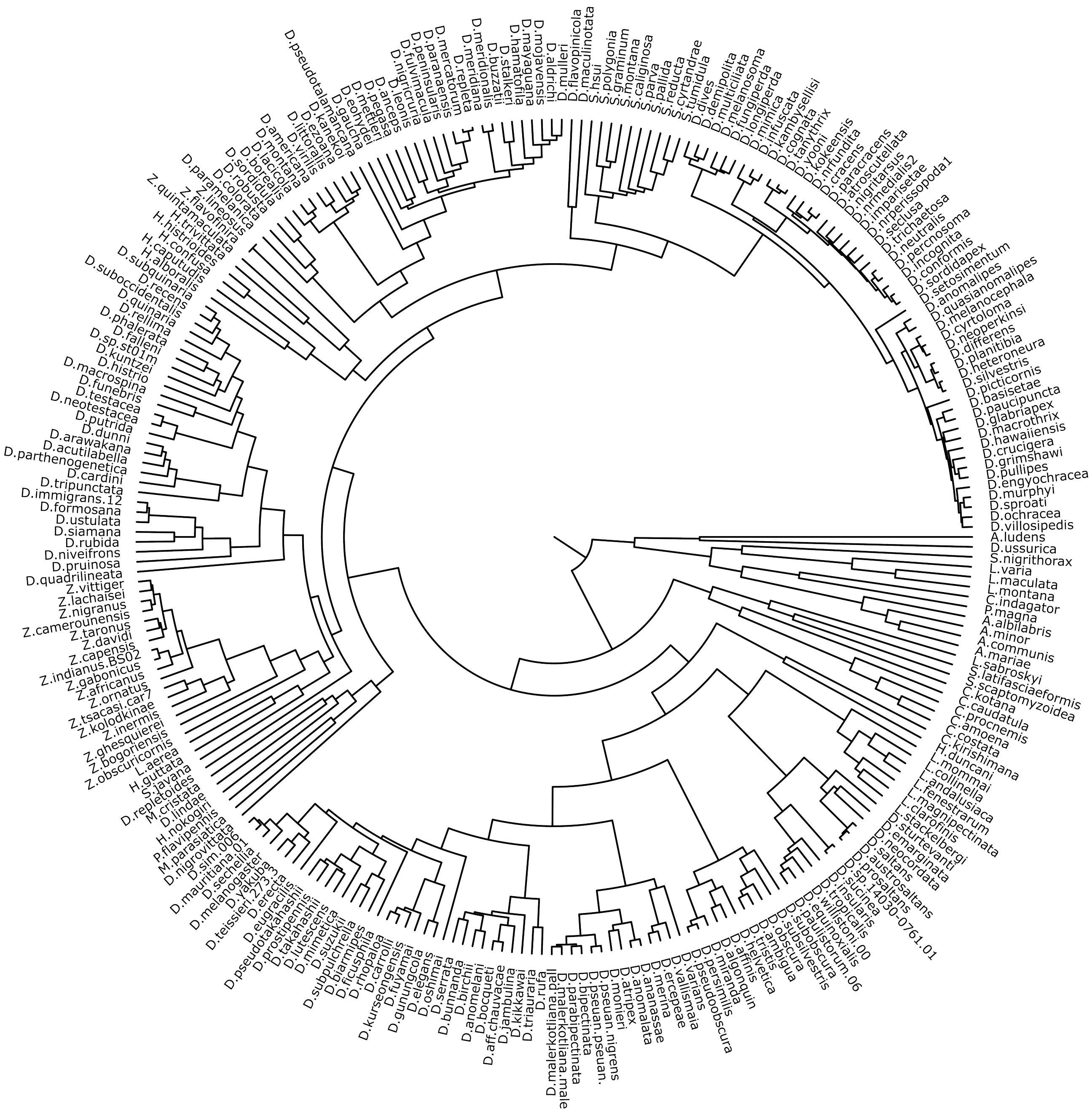


Figure 2: Bayesian phylogenetic tree of 262 drosophilids species investigated in this work. The tree was generated using orthologous genes (BUSCO).

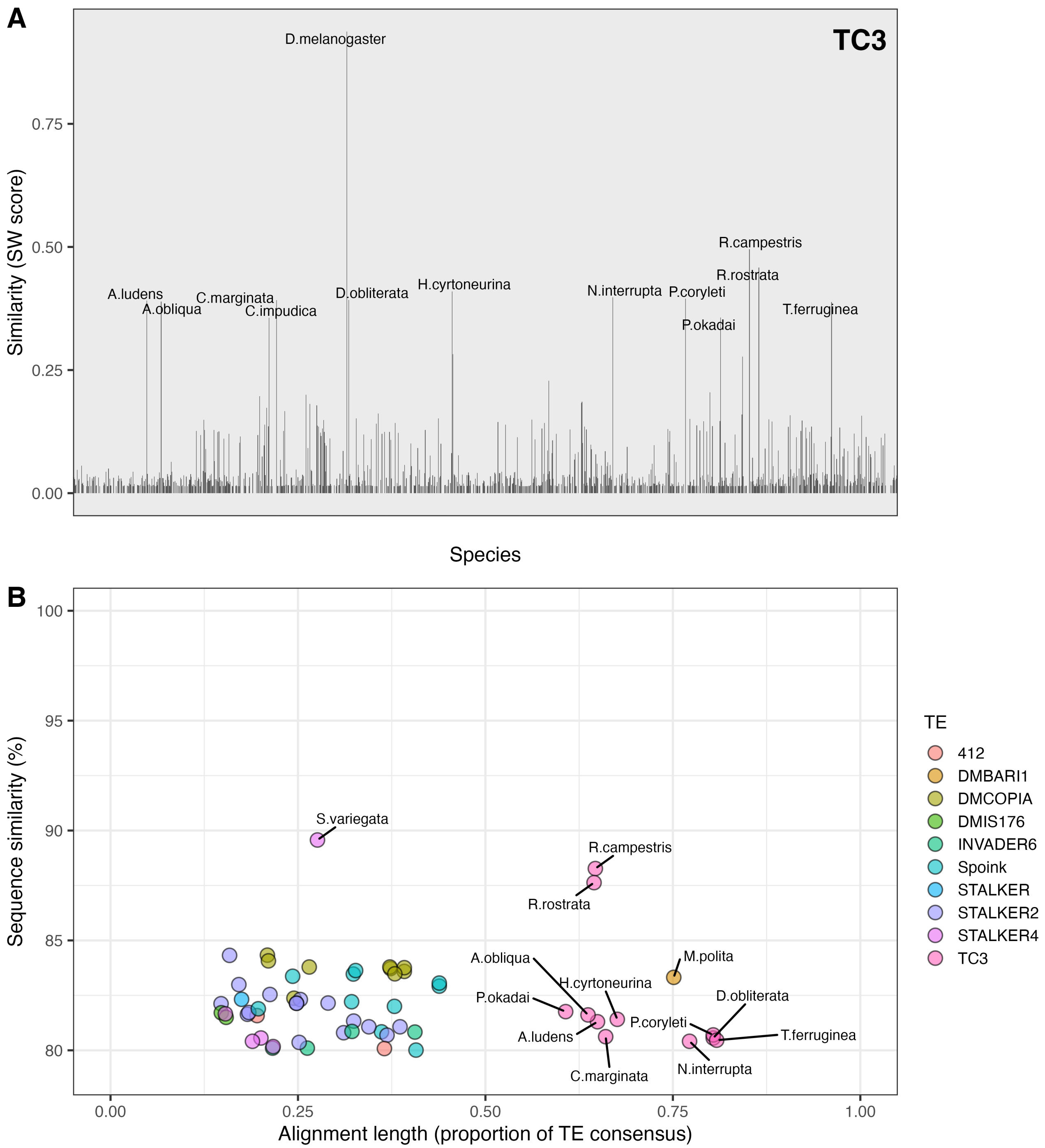


Figure 3: Sequences resembling *D. melanogaster* TEs in the genomes of 1200 arthropods (exluding drosophilids). A) Similarity of TC3 to sequences in the genomes of 1200 arthropods. Out of all investigated TEs, *D. melanogaster* TC3 has the most similar insertions in the 1200 arthropods. B) Summary of the sequences most closely resembling *D. melanogaster* TEs in arthropods. A recent HTT would lead to a high sequence similarity (close to 100%) and a high alignment length (close to 1.0). We did not observe any TE matching this criterion in arthropods, and thus conclude that *D. melanogaster* TEs were not involved in HTT with any of the investigated arthropod species.

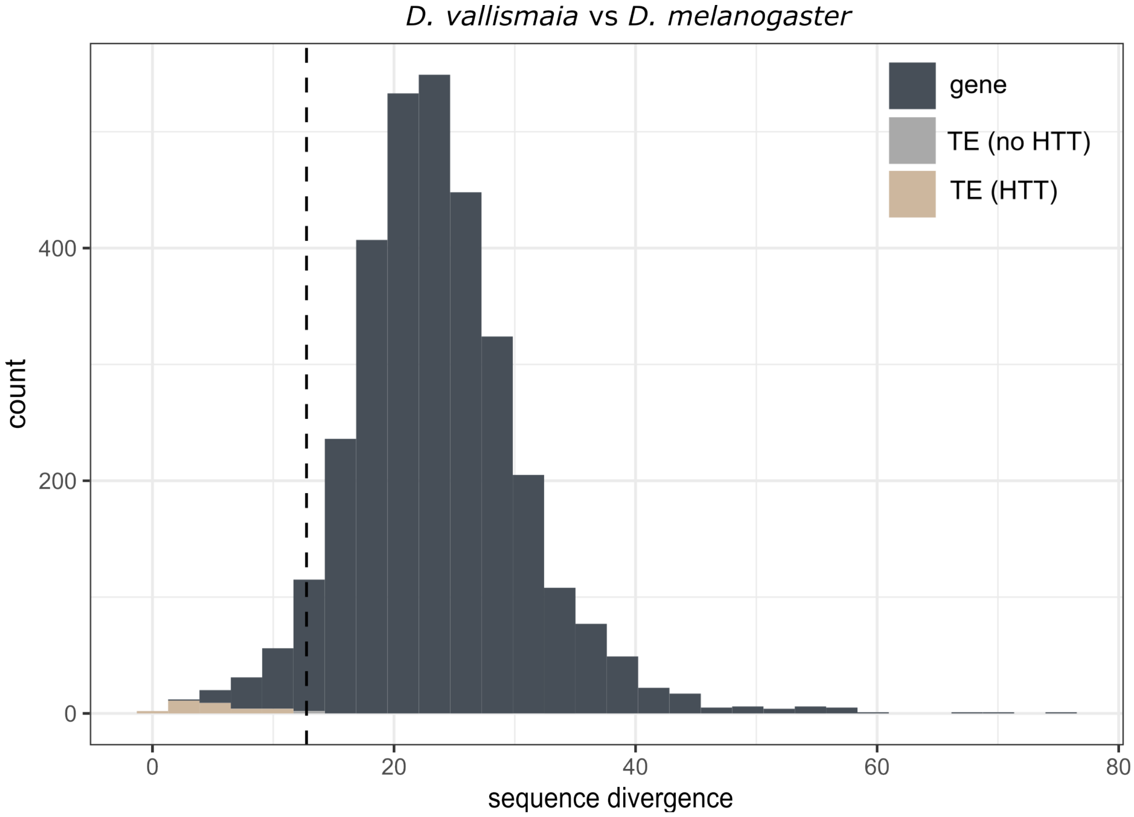


Figure 4: Overview of our test for detecting HTT. First, we computed the divergence of BUSCO genes between two species (e.g. *D. melanogaster* and *D.vallismaia*; dark grey). If the similarity of a TE among the two species is higher than 5% of the BUSCO genes (dashed lines) we assume a HTT (brown).

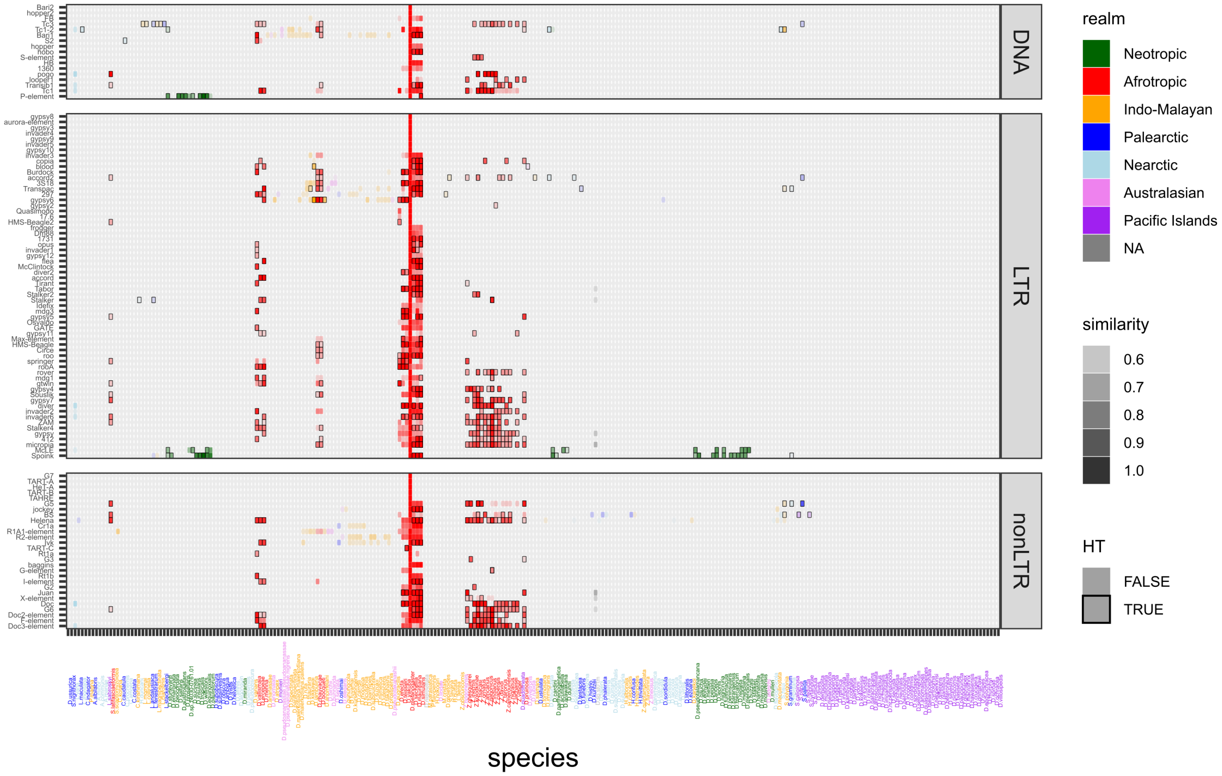


Figure 5: The matrix shows the similarity between *D. melanogaster* TEs (y-axis) and the best match in any of 262 drosophilid species (x-axis), separately for different orders of TEs. Intensity of the colors scales with the similarity score (panel on the right; based on Smith-Waterman score, which considers the sequence similarity as well as the length of a match). Only matches with a score > 0.5 are shown. Species were arranged according to their phylogeny and TEs according to the biogeographic origin. Black frames indicate likely HTT events with *D. melanogaster* as determined by our test.

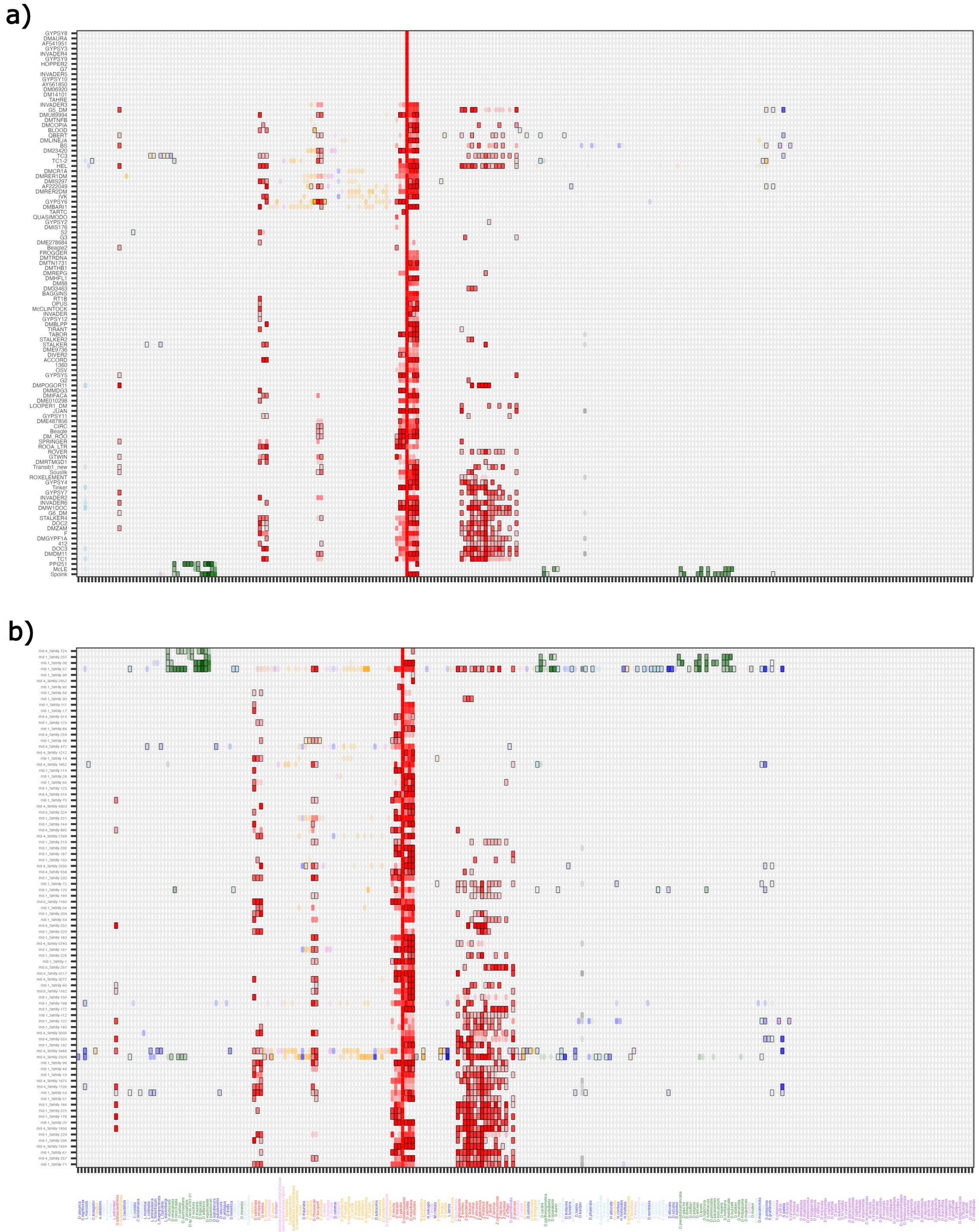


Figure 6: Distribution of sequences resembling *D. melanogaster* TEs in different drosophilid species using a high-quality manually curated library (A) and a repeat library generated by EarlGrey (B). The matrices show the similarity between *D. melanogaster* TEs (y-axis) and the best matching sequence in each of 262 drosophilid species (x-axis). The color reflects the biogeographic realm and the intensity of the colors scales with the similarity score (see text). Black frames indicate likely HTT events with *D. melanogaster* as determined by our test.

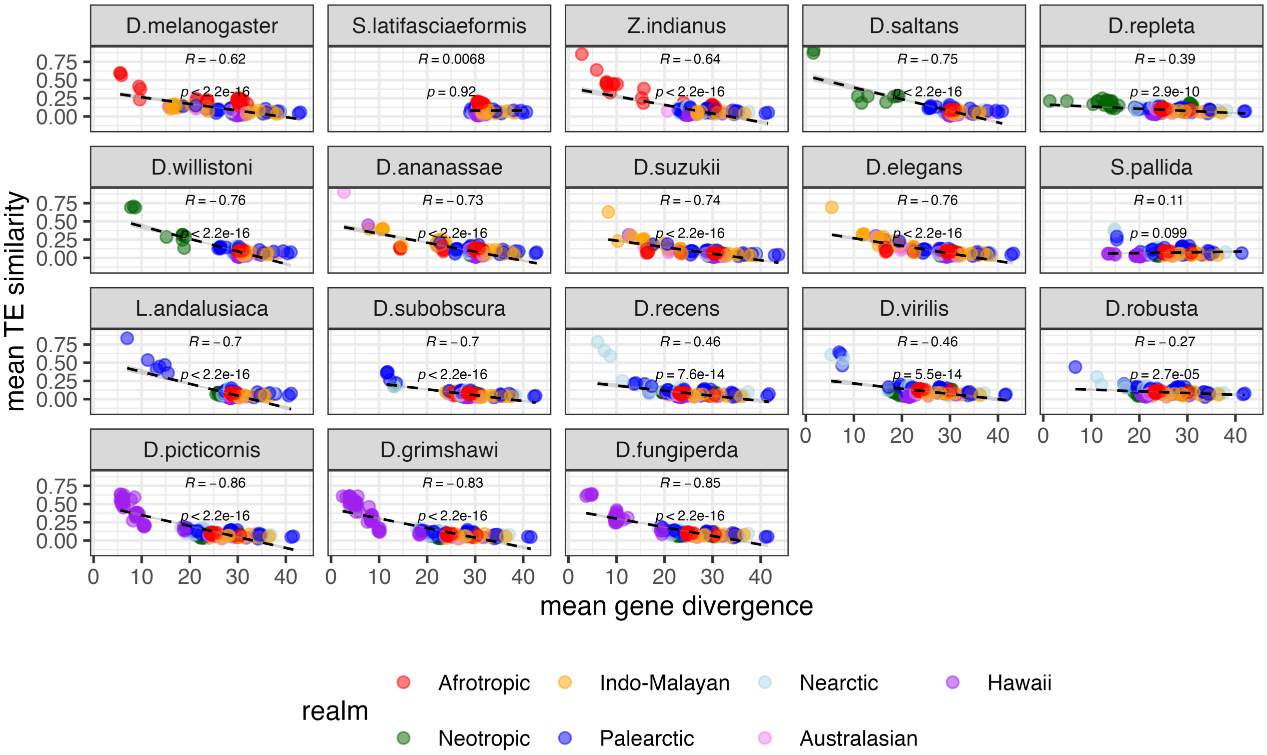


Figure 7: Scatter plots showing the average sequence similarity between TE of the focal species and TE sequences in 262 drosophilds (y-axis) versus the average divergence of BUSCO genes (x-axis). Spearman correlations are shown.

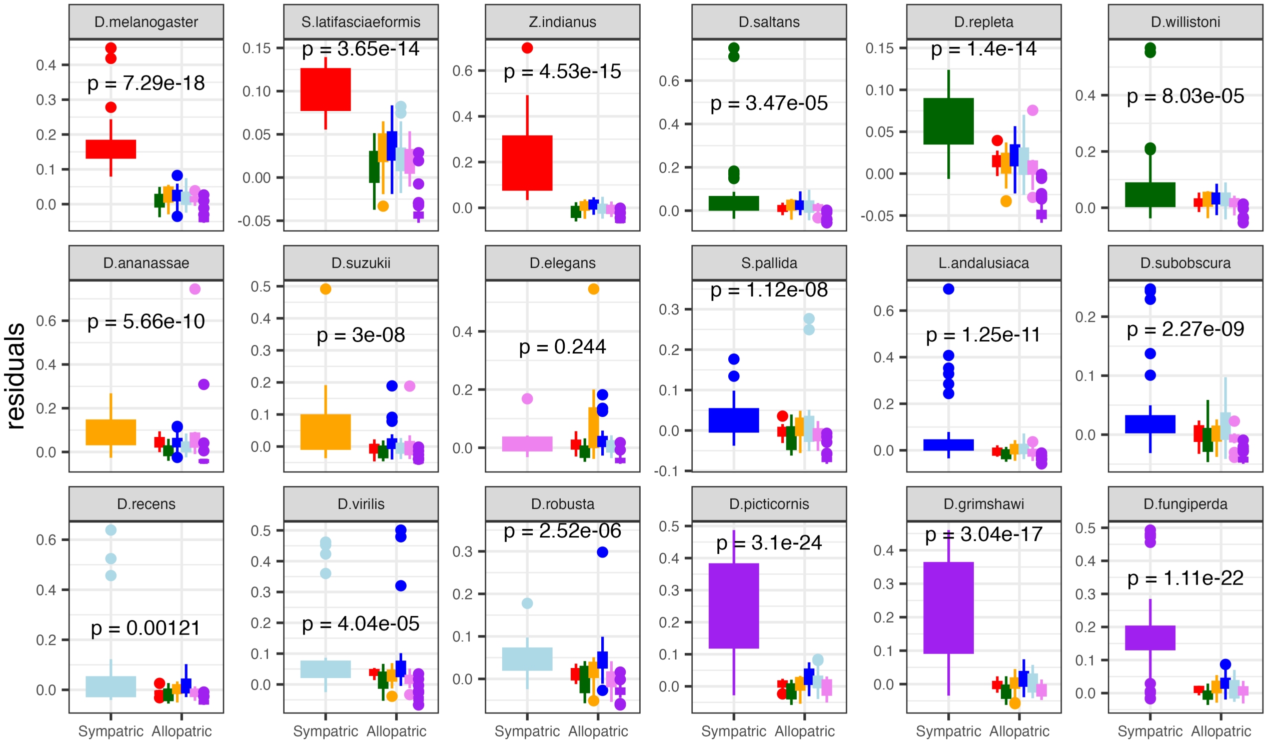


Figure 8: Residuals for 18 drosophilid species from diverse biogeographic realms (TEs were annotated with EarlGrey). Species from the same biogeographic realm as the focal species were not considered for generating the correlation, in contrast to a similar figure in the main manuscript where all species were used.

### Supplementary tables

Table 1: Genomic proportions due to TEs involved in HTT, TEs not involved in HTT (other TEs) and genomic sequences that were not annotated as TEs (non-mobile genome). Data are shown for 18 species from different biogeographic realms.

| **species** | **ancestral region** | **TEs involved in a HTT (%)** | **other TEs (%)** | **non-mobile genome (%)** |
|---:|---:|---:|---:|---:|
| *D. melanogaster* | Afrotropic | 14.99 | 2.07 | 82.94 |
| *S. latifasciaeformis* | Afrotropic | 15.30 | 3.79 | 80.91 |
| *Z. indianus* | Afrotropic | 32.35 | 25.68 | 41.97 |
| *D. ananassae* | EastAsia-Oceania | 28.23 | 7.47 | 64.30 |
| *D. suzukii* | EastAsia-Oceania | 55.09 | 3.00 | 41.91 |
| *D. elegans* | EastAsia-Oceania | 24.80 | 2.99 | 72.22 |
| *S. pallida* | Palearctic | 38.33 | 5.00 | 56.67 |
| *L. andalusiaca* | Palearctic | 32.31 | 7.37 | 60.32 |
| *D. subobscura* | Palearctic | 14.56 | 2.00 | 83.44 |
| *D. recens* | Nearctic | 21.09 | 1.94 | 76.97 |
| *D. virilis* | Nearctic | 23.60 | 3.14 | 73.26 |
| *D. robusta* | Nearctic | 10.71 | 3.32 | 85.97 |
| *D. willistoni* | Neotropic | 20.69 | 0.86 | 78.45 |
| *D. saltans* | Neotropic | 42.31 | 5.13 | 52.56 |
| *D. repleta* | Neotropic | 25.13 | 1.47 | 73.40 |
| *D. picticornis* | Hawaii | 4.52 | 0.88 | 94.60 |
| *D. grimshawi* | Hawaii | 22.17 | 21.83 | 56.00 |
| *D. fungiperda* | Hawaii | 3.25 | 0.77 | 95.98 |

### References

A. F. A. Smit, R. Hubley, and P. Green. RepeatMasker Open-4.0, 2013-2015. URL `http://www.repeatmasker.org`.
